# High CO_2_ promotes fatty acid-dependent asymbiotic growth of *Rhizophagus clarus*

**DOI:** 10.64898/2026.10.20.760566

**Authors:** Kayo Hashimoto, Sachiko Tanaka, Tatsuhiro Ezawa, Masayoshi Kawaguchi

**Affiliations:** Division of Symbiotic Systems, National Institute for Basic Biology, Nishigonaka 38, Myodaiji, Okazaki, Aichi 444-8585, Japan; Department of Basic Biology, School of Life Science, Graduate University for Advanced Studies (SOKENDAI), Nishigonaka 38, Myodaiji, Okazaki, Aichi 444-8585, Japan; Graduate School of Agriculture, Hokkaido University, Kita 9, Nishi 9, Kita-ku, Sapporo, Hokkaido, 060-8589, Japan

## Abstract

Arbuscular mycorrhizal (AM) fungi form symbioses with most terrestrial plants and play an ecologically important role in soil carbon dynamics. However, direct CO_2_ effects on the fungi remain unclear. We show that CO_2_-enriched conditions enhance the asymbiotic growth of *Rhizophagus clarus* in the presence of fatty acids. The response pattern to high CO_2_ differed between the tested species. These findings provide new insights into the physiological characteristics of AM fungi.

## Background

Arbuscular mycorrhizal (AM) fungi are obligate symbionts that provide plants with soil mineral nutrients in exchange for photosynthetically derived carbon^1^. Given the recent environmental changes at the global scale, the impact of elevated CO_2_ on AM symbiosis has attracted broad interest. Meta-analyses indicate that while these effects vary depending on factors such as AM fungal diversity and nitrogen availability, elevated CO_2_ generally enhances both AM colonisation and plant biomass^2^. In the pre-symbiotic stage, CO_2_ promotes hyphal development and is incorporated into fungal structures of *Gigaspora* sp. and *Rhizophagus* sp^3,4^. Recent studies showed that AM fungi rely on host-derived lipids for carbon^5,6,7,8^, as they lack essential fatty acid biosynthetic genes^9,10,11^. Some fatty acids induce asymbiotic spore formation in some *Rhizophagus* spp.^12,13^. GR24, a synthetic strigolactone, functions as a signalling molecule initiating AM symbiosis^14,15^, and accelerates asymbiotic growth in the presence of fatty acids, resulting in the mass production of spores in *R. clarus*^16^. These advances enable AM fungal responses to be examined without host plant involvement.

### Direct CO_2_ effects on *R. clarus*

To assess the direct CO_2_ effects on AM fungi, we modified the TG medium, which was developed based on the minimum (M) medium^17^ by adding myristate, peptone and GR24^16^. For this modified TG (mTG) medium, potassium dihydrogen phosphate and ammonium sulfate were added because the M medium contains very low concentrations of phosphate and nitrogen. Following previous reports^12,13^, potassium palmitate and potassium palmitoleate were supplied. The detailed medium composition is provided in Supplementary Table 1. Single spores of *R. clarus* were placed at the centre of the dishes and incubated under ambient, 0.5% and 2.0% CO_2_ conditions, representing typical concentrations in soil^18^. At 4 weeks after incubation (WAI), hyphae expanded radially with highly ramified branching, and many small spores were formed around parent spores under ambient and CO_2_-enriched conditions (Supplementary Fig. 1a). At 8 WAI, the fungi spread over the dishes and the supplied fatty acids that were visible as white precipitates in the medium were mostly consumed under CO_2_-enriched conditions (Fig. 1a and Supplementary Fig. 1a). *R. clarus* generally formed the largest spore clusters near the parent spore, as well as small clusters along runner hyphae regardless of conditions. Fungal biomass was evaluated as dry weight at 8 WAI. A growth-promoting effect, accompanied by the formation of numerous spores, was observed from 0.5% CO_2_ and became more pronounced at 2.0% (Fig. 1a,b). Relative biomass was higher under 2.0% CO_2_ than under 0.5% CO_2_ (Supplementary Fig. 1b). We further investigated higher CO_2_ concentrations (5.0–10.0%) reported under certain soil conditions^18^. Because the fungi proliferated across most of the dish at 6 WAI in about half of the 5.0% CO_2_ samples (Supplementary Fig. 1c), growth was evaluated at 6 WAI to avoid growth saturation. At 5.0% CO_2_, fungal biomass was consistently, but not significantly, higher than that at 2.0% across all trials. Conversely, fungal growth at 10.0% CO_2_ was significantly lower than at 2.0% and 5.0% (Supplementary Fig. 1c,d), indicating that 10.0% CO_2_ exceeded the optimal range for growth. The reduced or delayed growth at 10.0% CO_2_ may be related to its proximity to the upper range of reported soil CO_2_ concentrations^18^.

**Fig 1.**
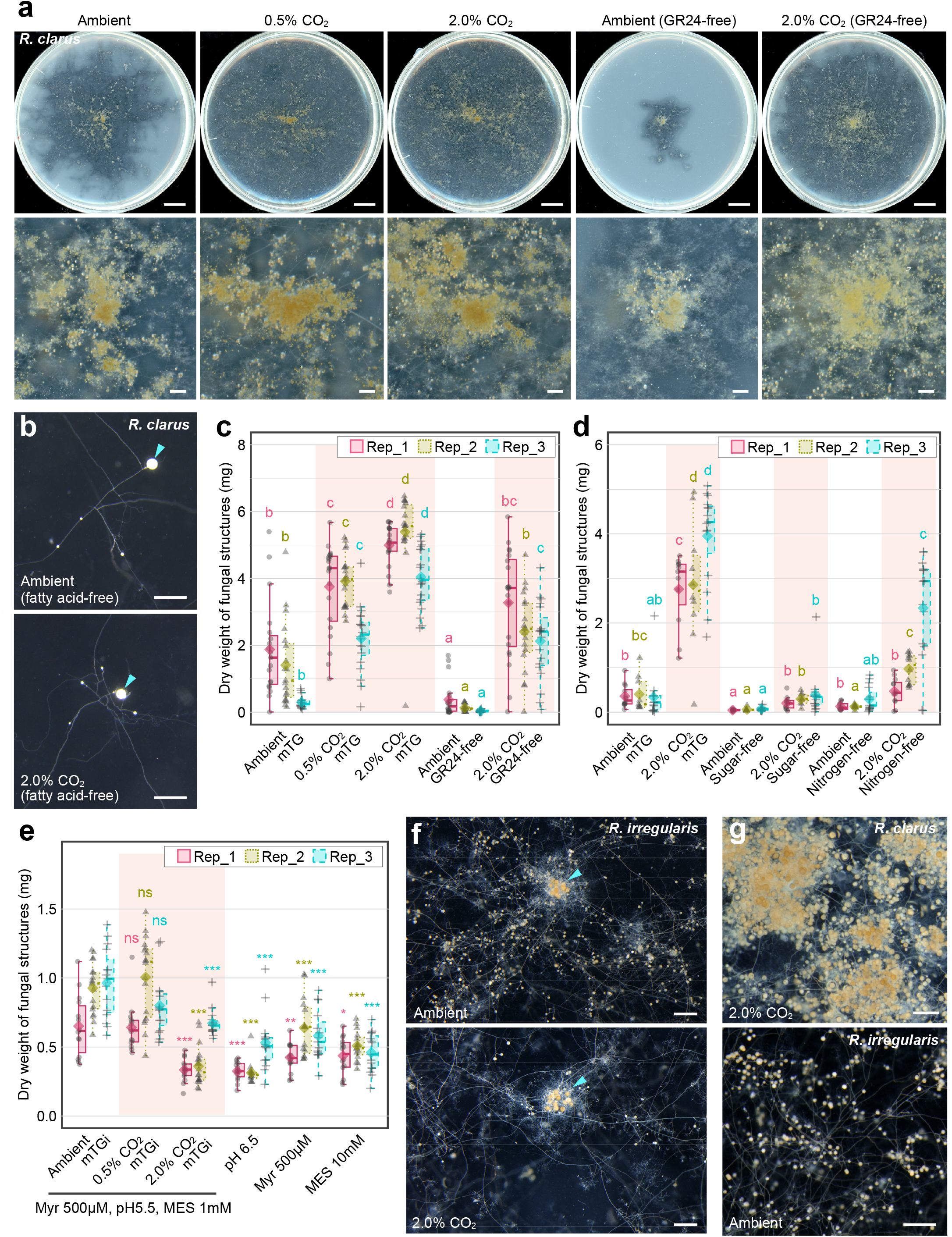
Asymbiotic growth in the medium containing fatty acids with or without GR24 under ambient or high concentration of CO_2_ conditions. **a**, Representative images of asymbiotically growing *R. clarus* in each condition. Top panels show whole dishes, and bottom panels show magnified images around parent spores. Bars, 1 cm (top) and 5 mm (bottom). **b**, Representative images of asymbiotically growing *R. clarus* in fatty acid-free media. Arrowheads represent parent spores. Bars, 500 µm. **c, d**, Dry weight of *R. clarus* under ambient, 0.5%, and 2.0% CO_2_ conditions in the presence or absence of GR24 (**c**) and under ambient or 2.0% CO_2_ conditions to investigate the requirement for sugars or nitrogen (**d**). Diamonds indicate means. Statistical significance was determined using the Steel–Dwass test (*p* < 0.05). Different letters indicate significant differences among conditions. *p*-values are described in Supplementary Table 2. **e**, Dry weight of fungal structures of *R. irregularis* under various atmospheric and medium conditions. Diamonds indicate means. Statistical significance to compare each treatment with the ambient mTGi control was determined using the Steel test (*p* < 0.05) with the mTGi medium under ambient conditions as a control. Asterisks indicate significant differences relative to the ambient mTGi control (***, *p* < 0.001; **, 0.001 ≤ *p* < 0.01; *, 0.01 ≤ *p* < 0.05). *p*-values are described in Supplementary Table 2. **f**, Representative images of asymbiotically growing *R. irregularis* in the mTGi medium under ambient and 2.0% CO_2_ conditions. Arrowheads represent 15 parent spores. Bars, 500 µm. **g**, Comparison of spore-forming patterns between *R. clarus* and *R. irregularis* under favourable atmospheric conditions for each fungus. Bars, 500 µm.

### Medium components contributing to CO_2_-induced growth promotion

The effects of fatty acids—potassium myristate, palmitate, and palmitoleate—on fungal growth varied depending on their concentrations and CO_2_ levels (Supplementary Fig. 1f–j). Under ambient conditions, biomass remained largely unchanged across most fatty acid combinations, except for a higher concentration of potassium palmitoleate, which severely suppressed growth. Nonetheless, high CO_2_ significantly promoted fungal growth across all fatty acid combinations. Under high CO_2_, biomass tended to increase with higher concentrations of potassium myristate, whereas growth inhibition induced by potassium palmitoleate was alleviated. These observations imply that CO_2_-enriched conditions enhance the efficiency of fatty acid utilisation in *R. clarus*. Crucially, *R. clarus* displayed stunted growth with limited hyphal branching and only a few small spontaneous spores in fatty acid-free medium under both ambient and 2.0% CO_2_ conditions (Fig. 1b and Supplementary Fig. 1e). This demonstrates that fatty acids are essential for CO_2_-driven growth promotion.

Strigolactone activates cellular metabolism in AM fungi^15,19^, while myristate enhances metabolic pathways^13^. Although both have similar stimulatory effects, they act synergistically on asymbiotic growth^16^. Consistent with this, GR24 significantly promoted *R. clarus* growth under both atmospheric conditions. Both CO_2_ and GR24 exerted independent effects on fungal biomass without significant interaction between them (Fig. 1c and Supplementary Table 3). However, the relative biomass increase induced by GR24 was markedly smaller under 2.0% CO_2_ than under ambient conditions. Under 2.0% CO_2_, biomass in the GR24-free medium reached 35-fold that under ambient conditions (Supplementary Fig. 1b). While fungal growth in the mTG medium under ambient conditions was occasionally limited depending on the trial, high CO_2_ was consistently associated with higher growth even in the absence of GR24.

In sugar- or nitrogen-free media, high CO_2_ promoted fungal growth, and both CO_2_ and nutrients had independent effects on biomass without significant interaction (Fig. 1d, Supplementary Fig. 1k and Supplementary Table 3). CO_2_-induced growth promotion was lower in the absence of sugars or nitrogen than in the mTG medium (Supplementary Fig. 1l), indicating that these nutrients partially contribute to the growth promotion. Similar to myristate, and in contrast to GR24, the relative effect of each nutrient was greater under 2.0% CO_2_ than under ambient conditions, suggesting that a CO_2_-enriched environment enhances the utilisation efficiency of these nutrients, particularly sugars.

### CO_2_ responses in other AM fungal species

To date, AM fungi capable of asymbiotic propagation on fatty acid-supplemented media have been limited to specific taxa, such as *Rhizophagus*^12,13,16,20^. In *G. rosea*, no new spores were produced even in the presence of both high CO_2_ and myristate, although a few auxiliary cells were formed (Supplementary Fig. 1m). *R. irregularis* exhibited substantially less consistent growth upon single-spore inoculation than *R. clarus*^16^. To facilitate asymbiotic growth of *R. irregularis*, 15 spores were placed in a cluster. The medium composition was also optimised by using a more acidic pH (5.5) and lower concentration of myristate (200 µM) and MES (1 mM; Supplementary Table 1) than those used in the mTG medium. These modifications successfully reduced inter-sample variability, facilitating a clearer observation of growth trends. On this medium (mTG for *R. irregularis*; mTGi), 0.5% CO_2_ had no significant effect on biomass, unlike in *R. clarus*. Notably, in contrast to *R. clarus*, 2.0% CO_2_ had an inhibitory effect (Fig. 1e,f and Supplementary Fig. 1n).

Compared to *R. clarus, R. irregularis* generally developed extensive runner hyphae with sparse spores formed in distant areas rather than around parent spores (Fig. 1g and Supplementary Fig. 1n,o). This tendency observed in the mTGi medium regardless of atmospheric conditions. Conversely, on media with high concentration of myristate or MES, *R. irregularis* tended to form spores primarily around parent spores, similar to *R. clarus*, although colony expansion was limited and biomass was lower (Supplementary Fig. 1o). Recent studies have demonstrated that while AM fungi can adjust their behaviour under different nutrient conditions, their strategies are constrained within species-specific ranges^21^. In this framework, *R. irregularis* tends toward an “explorative” strategy—prioritising rapid expansion—whereas *R. clarus* is an intermediate type with a more “exploitative” tendency, producing a relatively higher hyphal density. This appears to align with qualitative differences between species and plastic responses to media conditions observed in asymbiotic cultures. Quantitative studies across more fungal isolates will be required to elucidate their trait diversity.

### Transcriptomic analysis

To investigate the mechanism behind CO_2_-induced growth promotion in *R. clarus*, RNA-Seq was conducted. Reduced-gellan gum medium^22^ was employed because recovering fungal structures from solid agar medium is time-consuming. Principal component and hierarchical clustering analyses separated gene expression profiles between ambient and 2.0% CO_2_ conditions (Supplementary Fig. 2). Of 30,194 predicted protein-coding genes, 16,447 remained after filtering. Among these, 973 and 939 were identified as up- and down-regulated differentially expressed genes (DEGs), respectively (Supplementary Fig. 2 and Supplementary Data 1). In the GO enrichment analysis, up-regulated DEGs were enriched in terms related to protein modification and signal transduction, and down-regulated DEGs were enriched in those related to transport and structural organisation (Supplementary Fig. 2).

### Perspectives on CO_2_ utilization

CO_2_ affects fungal growth and morphogenesis through multiple mechanisms, such as carbon assimilation, activation of cAMP signalling^23^, and the modulation mitochondrial respiration and redox balance^24^. The remarkable stimulation of AM fungal growth under 0.5– 5.0% CO_2_ likely reflects physiological adaptation of AM fungi to the soil environments, where CO_2_ concentrations are elevated by root and microbial respiration and restricted gas diffusion^18^. However, our RNA-Seq identified no GO terms directly associated with these processes. Physiological heterogeneity likely diluted stage-specific signals and masked localised transcriptional responses to CO_2_-enriched conditions. Future analysis using stage- or tissue-specific samples, together with metabolomic and comparative genomic approaches, may help elucidate the molecular mechanisms underlying CO_2_ utilisation in AM fungi. Given the importance of AM fungi in soil carbon storage^25^, extending our laboratory findings across fungal species and environmental conditions could advance our understanding of the fungal traits and their potential relevance to sustainable agriculture.

## Methods

### Fungal materials and asymbiotic cultures

Fungal strains of *R. clarus* HR1 (MAFF520076) and *G. rosea* (MAFF520062) were obtained from NARO Genebank and *R. irregularis* (DAOM197198) was purchased from Premier Tech. To be used as inoculum, spores of *R. clarus* and *R. irregularis* were cultivated by monoxenic culture system with carrot hairy roots for at least 3 months. *G. rosea* spores were collected from soil cultures with *Lotus japonicus* after 4 months of cultivation, followed by 1 month of drying and subsequent storage at 4°C. Medium preparation and fungal inoculation were performed according to previous reported methods^16,26^. For *G. rosea*, the manganese concentration was reduced to prevent spore germination inhibition^27^. The detailed medium composition is described in Supplementary Table 1. Sugar- and nitrogen-free media were prepared by omitting glucose/sucrose and nitrate/ammonium/organic nitrogen, respectively. Culture media solidified with 2% gellan gum (w/v) were poured into 9 cm Petri dishes (20 ml for *R. clarus* and 15 ml for *R. irregularis*). For inoculation onto the center of each dish, a single spore exceeding 140 µm in diameter was used for *R. clarus*, and 15 spores exceeding 80 µm were used for *R. irregularis*. The fungi were grown at 28°C in the dark. Ambient conditions were established using an LH411S incubator (Nippon Medical & Chemical Instruments Co., Ltd., Japan), whereas CO-enriched conditions were created using MCO-18AIC, MCO-19AIC and MCO-170AIC incubators (PHC, Japan). To allow gas exchange, dishes were wrapped with a 10 cm-long strip of Parafilm while stretching the film. All experiments were repeated three times, using spores derived from different monoxenic culture dishes for each independent trial. For fungal growth evaluation, ungerminated spores and those that showed stunted growth with only a few branched hyphae and no spores were excluded (Supplementary Table 4). Fungal dry weight was measured according to a protocol book^26^ using MX5 micro analytical balance (METTLER TOLEDO, USA). Fungal structures were observed under an SZX7 stereomicroscope (Olympus, Japan) or with a GT-X980 flatbed scanner (EPSON, Japan).

### RNA-Seq

#### Sample preparation, RNA extraction, library preparation, and sequencing

*R. clarus* was cultured in the mTG medium with 0.04% gellan gum under ambient or 2.0% CO_2_ conditions^22^. For ambient conditions, three spores were inoculated per dish. For 2.0% CO_2_ conditions, single spores were inoculated per dish to avoid colony overlap due to enhanced growth. In total, 9–10 and 12–14 dishes, respectively, were prepared across for each three trials. Fungal structures were harvested at 4 WAI to obtain samples during active growth. The sampling procedure was as follows: culture dishes were shaken by hand for 2 s, and the resulting slurry containing fungal structures was immediately poured into 150 mL of citrate buffer (55°C) and swirled for a few seconds. After confirming the complete dissolution of the gel blocks, the fungal suspension was filtered through a funnel with 22 µm nylon mesh. The mesh was washed thoroughly with 300 mL of distilled water and placed on filter paper to remove excess moisture. Fungal structures were collected using forceps or scrapers, transferred into sampling tubes, and immediately frozen in liquid nitrogen. Total RNA extraction, library preparation, and RNA-Seq were performed by Bioengineering Lab. Co., Ltd. (Japan). Fungal structures were disrupted using a Multi-beads Shocker (Yasui Kikai, Japan), and total RNA was extracted using the RNeasy Mini Kit on a QIAcube Connect system (Qiagen, Netherlands). RNA libraries were constructed using the MGIEasy Fast RNA Library Prep Set (MGI Tech, China). The libraries were circularised with the MGIEasy Dual Barcode Circularisation Kit and converted into DNA nanoballs using the DNB Rapid Make Reagent Kit (MGI Tech). Sequencing was conducted on the DNBSEQ-T7 platform (MGI Tech) with 150-bp paired-end reads.

### Pre-processing and quality control

Adapter sequences were removed using Cutadapt (v.5.1), and low-quality (Q ≤ 20) or short (≤ 20 bp) reads were trimmed using Sickle (v.1.33). Read quality was assessed using FastQC (v.0.12.1). To remove ribosomal RNA (rRNA) contaminants, reads were mapped to 11 rRNA sequences identified in the *R. clarus* genome (NCBI accession: GCA_015698045.1) using Bowtie2 (v.2.4.4). The resulting unmapped reads were extracted and utilised for downstream analyses.

### Transcript quantification and differential expression analysis

Transcript abundance was quantified using Salmon (v.1.10.3) with the rRNA-filtered reads obtained from the Bowtie2 filtering step. The transcriptome sequence was extracted from the GFF file using gffread (v.0.12.7) and used as a reference to build an index for Salmon. Differential expression analysis was performed using the TCC-GUI pipeline^28^. Raw counts were normalised using the trimmed mean of M-values method, and DEGs were identified using edgeR (v.4.6.3). DEGs were defined based on the criteria of a false discovery rate (FDR) < 0.05, with *p*-value thresholds set at 0.05.

### Functional annotation and enrichment analysis

Functional annotations were assigned to the protein sequences (protein.faa from NCBI accession: GCA_015698045.1) using InterProScan (5.75-106.0). GO enrichment analysis was performed using the enricher function in the R package clusterProfiler (v.4.16.0), and results were visualised using the enrichplot package (v.1.28.4). DEGs were categorised into upregulated (M **≥** 1) and downregulated (M ≤ −1) sets for separate analysis. Enrichment analysis used Benjamini– Hochberg adjustment method, a *p*-value cutoff of 0.05, an FDR cutoff of 0.05, and a minimum gene set size of 5.

### Statistical analysis

Statistical analyses for fungal dry weight and colony size were performed using R (v.4.5.0) as follows. The Wilcoxon rank-sum test was used for comparisons between two groups. For multiple comparisons, the Steel test was used to compare each treatment group against the control group, while the Steel–Dwass test was applied for all-pairwise comparisons among all groups with the nparcomp package (v.3.0). To evaluate the effects of atmospheric and medium conditions on fungal growth, the Scheirer–Ray–Hare test was performed with the rcompanion package (v. 2.5.2). *p*-values and H statistics are presented in Supplementary Table 3.

## Data availability

All data used for culture experiments are included in the main or supplementary information. The raw sequence data generated in this study have been deposited in the DDBJ Sequence Read Archive under the BioProject PRJDB40706.

## Acknowledgements

We are grateful to Koji Yano, Takumi Sato, Shingo Aoki, Mikio Fujii, and Shintaro Suzuki (Nagase ChemteX Co., Ltd.) for their suggestions regarding the experimental design for evaluating the effects of different gas conditions on fungal growth using AnaeroPack (Mitsubishi Gas Chemical Company, Inc., Japan). We thank Akiko Oda for experimental support, Kagayaki Kato for advice on statistical treatment, and the Model Organisms and the Trans-Omics Facilities for providing equipment (all at NIBB).

## Author contributions

MK and KH conceived and designed the experiments. KH performed all experiments, conducted the data analysis, and wrote the original draft of the manuscript. ST provided technical support for the experiments. TE contributed to the initial conception of the study and provided expert knowledge for the revision of the manuscript. MK supervised the study.

## References

1. Smith, S. E. & Read, D. Mycorrhizal Symbiosis (Academic Press, 2008).

2. Dong, Y., Wang, Z., Sun, H., Yang, W. & Xu, H. The Response Patterns of Arbuscular Mycorrhizal and Ectomycorrhizal Symbionts Under Elevated CO2: A Meta-Analysis. Front Microbiol. 9, 1248 (2018).

3. Becard, G. & Piche, Y. Fungal Growth Stimulation by CO2 and Root Exudates in Vesicular-Arbuscular Mycorrhizal Symbiosis. Appl Environ Microbiol. 55, 2320–2325 (1989).

4. Bago, B. et al. Carbon metabolism in spores of the arbuscular mycorrhizal fungus Glomus intraradices as revealed by nuclear magnetic resonance spectroscopy. Plant Physiol. 121, 263–272 (1999).

5. Kameoka, H. & Gutjahr, C. Functions of Lipids in Development and Reproduction of Arbuscular Mycorrhizal Fungi. Plant Cell Physiol. 63, 1356–1365 (2022).

6. Jiang, Y. et al. Plants transfer lipids to sustain colonization by mutualistic mycorrhizal and parasitic fungi. Science. 356, 1172–1175 (2017).

7. Luginbuehl, L. H. et al. Fatty acids in arbuscular mycorrhizal fungi are synthesized by the host plant. Science. 356, 1175–1178 (2017).

8. Keymer, A. et al. Lipid transfer from plants to arbuscular mycorrhiza fungi. Elife. 6 (2017).

9. Wewer, V., Brands, M. & Dormann, P. Fatty acid synthesis and lipid metabolism in the obligate biotrophic fungus Rhizophagus irregularis during mycorrhization of Lotus japonicus. Plant J. 79, 398–412 (2014).

10. Tisserant, E. et al. Genome of an arbuscular mycorrhizal fungus provides insight into the oldest plant symbiosis. Proc Natl Acad Sci U S A. 110, 20117–20122 (2013).

11. Kobayashi, Y. et al. The genome of Rhizophagus clarus HR1 reveals a common genetic basis for auxotrophy among arbuscular mycorrhizal fungi. BMC Genomics. 19, 465 (2018).

12. Kameoka, H. et al. Stimulation of asymbiotic sporulation in arbuscular mycorrhizal fungi by fatty acids. Nat Microbiol. 4, 1654–1660 (2019).

13. Sugiura, Y. et al. Myristate can be used as a carbon and energy source for the asymbiotic growth of arbuscular mycorrhizal fungi. Proc Natl Acad Sci U S A. 117, 25779–25788 (2020).

14. Akiyama, K., Matsuzaki, K. & Hayashi, H. Plant sesquiterpenes induce hyphal branching in arbuscular mycorrhizal fungi. Nature. 435, 824–827 (2005).

15. Besserer, A. et al. Strigolactones stimulate arbuscular mycorrhizal fungi by activating mitochondria. PLoS biology. 4, e226 (2006).

16. Tanaka, S. et al. Asymbiotic mass production of the arbuscular mycorrhizal fungus Rhizophagus clarus. Commun Biol. 5, 43 (2022).

17. Becard, G. & Fortin, J. A. Early events of vesicular-arbuscular mycorrhiza formation on Ri T-DNA transformed roots. New Phytologist. 108, 211–218 (1988).

18. Amundson, R. G. & Davidson, E. A. Carbon dioxide and nitrogenous gases in the soil atmosphere. Journal of Geochemical Exploration. 38, 13–41 (1990).

19. Besserer, A., Becard, G., Jauneau, A., Roux, C. & Sejalon-Delmas, N. GR24, a synthetic analog of strigolactones, stimulates the mitosis and growth of the arbuscular mycorrhizal fungus Gigaspora rosea by boosting its energy metabolism. Plant Physiol. 148, 402–413 (2008).

20. Cha, J. E. & Eom, A. H. Asymbiotic Spore Production of Rhizoglomus intraradices in a Medium Containing Myristate. Mycobiology. 51, 164–168 (2023).

21. Bisot, C. et al. Carbon-phosphorus exchange rate constrains density-speed trade-off in arbuscular mycorrhizal fungal growth. Proc Natl Acad Sci U S A. 123, e2512182123 (2026).

22. Kawaguchi, M. et al. MEDIUM AND METHOD FOR CULTURING ARBUSCULAR MYCORRHIZAL FUNGUS, 2020).

23. Chadwick, B. J. & Lin, X. Effects of CO2 in fungi. Curr Opin Microbiol. 79, 102488 (2024).

24. Ristow, L. C., Blackburn, E. E., Jezewski, A. J., Lin, X. & Krysan, D. J. Cryptococcus neoformans adapts to host CO2 concentrations via metabolic and stress-response remodeling. PLoS biology. 24, e3003561 (2026).

25. Hawkins, H. J. et al. Mycorrhizal mycelium as a global carbon pool. Curr Biol. 33, R560–R573 (2023).

26. Hashimoto, K., Tanaka, S. & Kawaguchi, M. Asymbiotic growth of arbuscular mycorrhizal fungi. in Arbuscular Mycorrhizal Fungi (eds Ferrol, N. & Lanfranco, L.) 19–32 (Humana Press, 2026).

27. Mizonobe, G. Some Conditions of Hyphae Germination from Sterilized Spores of Gigaspora margarita. Journal of Hokkaido Junior College, Senshu University, 1–8 (1999).

28. Su, W., Sun, J., Shimizu, K. & Kadota, K. TCC-GUI: a Shiny-based application for differential expression analysis of RNA-Seq count data. BMC Res Notes. 12, 133 (2019).

