## Supplementary Information for "High CO_2_ promotes fatty acid-dependent asymbiotic growth of *Rhizophagus clarus*"

**Supplementary Table 1.** Compositions and final concentrations of base media for asymbiotic culture experiments

<table>
<tr><th>Chemicals</th><th>TG medium[16]</th><th>mTG medium<br>optimized for<br>R. clarus</th><th>mTGi medium<br>optimized for<br>R. irregularis</th><th>Medium<br>used for<br>G. rosea</th></tr>
<tr><td>$MgSO_4 \cdot 7H_2O$</td><td>731 mg $l^{-1}$</td><td colspan="2" rowspan="11">Same as the TG medium</td><td rowspan="5">Same as the TG medium</td></tr>
<tr><td>$KNO_3$</td><td>80 mg $l^{-1}$</td></tr>
<tr><td>KCl</td><td>65 mg $l^{-1}$</td></tr>
<tr><td>$Ca(NO_3)_2 \cdot 4H_2O$</td><td>288 mg $l^{-1}$</td></tr>
<tr><td>Fe(III)-EDTA</td><td>8 mg $l^{-1}$</td></tr>
<tr><td>$MnCl_2 \cdot 4H_2O$</td><td>3 mg $l^{-1}$</td><td>0.15 mg $l^{-1}$</td></tr>
<tr><td>$ZnSO_4 \cdot 7H_2O$</td><td>1.3 mg $l^{-1}$</td><td rowspan="5">Same as the TG medium</td></tr>
<tr><td>$H_3BO_3$</td><td>1.5 mg $l^{-1}$</td></tr>
<tr><td>$CuSO_4 \cdot 5H_2O$</td><td>0.065 mg $l^{-1}$</td></tr>
<tr><td>$Na_2MoO_4 \cdot 2H_2O$</td><td>0.0012 mg $l^{-1}$</td></tr>
<tr><td>KI</td><td>0.75 mg $l^{-1}$</td></tr>
<tr><td>$KH_2PO_4$</td><td>4.8 mg $l^{-1}$</td><td>204.8 mg $l^{-1}$</td><td>204.8 mg $l^{-1}$</td><td>4.8 mg $l^{-1}$</td></tr>
<tr><td>$(NH_4)_2SO_4$</td><td>-</td><td>200 mg $l^{-1}$</td><td>200 mg $l^{-1}$</td><td>200 mg $l^{-1}$</td></tr>
<tr><td>Sucrose</td><td>1 g $l^{-1}$</td><td colspan="2" rowspan="9">Same as the TG medium</td><td rowspan="8">Same as the TG medium</td></tr>
<tr><td>Glucose</td><td>1 g $l^{-1}$</td></tr>
<tr><td>Glycine</td><td>3 mg $l^{-1}$</td></tr>
<tr><td>Pyridoxine-HCl</td><td>0.1 mg $l^{-1}$</td></tr>
<tr><td>Nicotinic acid</td><td>0.5 mg $l^{-1}$</td></tr>
<tr><td>myo-inositol</td><td>50 mg $l^{-1}$</td></tr>
<tr><td>Thiamine-HCl</td><td>10 mg $l^{-1}$</td></tr>
<tr><td>GR24</td><td>100 nM</td></tr>
<tr><td>Peptone</td><td>1 mg $l^{-1}$</td><td>0.5 mg $l^{-1}$</td></tr>
<tr><td>MES buffer</td><td>pH 6.5, 10 mM</td><td>pH 6.5, 10 mM</td><td>pH 5.5, 1 mM</td><td>pH 6.0, 1 mM</td></tr>
<tr><td>Potassium myristate</td><td>500 µM</td><td>500 µM</td><td>200 µM</td><td>100 µM</td></tr>
<tr><td>Potassium palmitate</td><td>-</td><td>200 µM</td><td>80 µM</td><td>20 µM</td></tr>
<tr><td>Potassium palmitoleate</td><td>-</td><td>0.1 µM</td><td>1 µM</td><td>-</td></tr>
</table>

**Supplementary Table 2.** Medians and fold changes (FC) and geometric means of fungal biomass under various culture conditions. The conditions used for the normalization are underlined. Daggers (†) indicate the same sample; the control mTG medium.

**Fig. 1c and Supplementary Fig. 1b**

| | Median | | | FC (relative to Ambient) | | | **Geometric mean (FC)** |
|---|---|---|---|---|---|---|---|
| Condition | Rep_1 | Rep_2 | Rep_3 | Rep_1 | Rep_2 | Rep_3 | |
| Ambient, mTG | 1.63 | 0.95 | 0.24 | 1.00 | 1.00 | 1.00 | **1.00** |
| 0.5% $CO_2$, mTG | 4.31 | 3.97 | 2.32 | 2.64 | 4.19 | 9.58 | **4.73** |
| 2.0% $CO_2$, mTG | 5.07 | 5.57 | 3.96 | 3.11 | 5.87 | 16.33 | **6.68** |
| Ambient, GR24-free | 0.17 | 0.11 | 0.03 | 0.11 | 0.11 | 0.12 | **0.11** |
| 2.0% $CO_2$, GR24-free | 3.71 | 2.50 | 2.41 | 2.27 | 2.63 | 9.95 | **3.91** |

| | Median | | | FC (relative to Ambient) | | | **Geometric mean (FC)** |
|---|---|---|---|---|---|---|---|
| Condition | Rep_1 | Rep_2 | Rep_3 | Rep_1 | Rep_2 | Rep_3 | |
| Ambient, GR24-free | 0.17 | 0.11 | 0.03 | 1.00 | 1.00 | 1.00 | **1.00** |
| 2.0% $CO_2$, GR24-free | 3.71 | 2.50 | 2.41 | 21.66 | 23.39 | 84.58 | **34.99** |

| | Median | | | FC (relative to GR24-free) | | | **Geometric mean (FC)** |
|---|---|---|---|---|---|---|---|
| Condition | Rep_1 | Rep_2 | Rep_3 | Rep_1 | Rep_2 | Rep_3 | |
| Ambient, GR24-free | 0.17 | 0.11 | 0.03 | 1.00 | 1.00 | 1.00 | **1.00** |
| Ambient, mTG | 1.63 | 0.95 | 0.24 | 9.52 | 8.89 | 8.50 | **8.95** |
| 2.0% $CO_2$, GR24-free | 3.71 | 2.50 | 2.41 | 1.00 | 1.00 | 1.00 | **1.00** |
| 2.0% $CO_2$, mTG | 5.07 | 5.57 | 3.96 | 1.37 | 2.23 | 1.64 | **1.71** |

**Supplementary Fig. 1d**

| | Median | | | FC (relative to 2.0% $CO_2$) | | | **Geometric mean (FC)** |
|---|---|---|---|---|---|---|---|
| Condition | Rep_1 | Rep_2 | Rep_3 | Rep_1 | Rep_2 | Rep_3 | |
| 2.0% $CO_2$, mTG | 1.66 | 2.52 | 2.76 | 1.00 | 1.00 | 1.00 | **1.00** |
| 5.0% $CO_2$, mTG | 4.15 | 3.55 | 3.74 | 2.49 | 1.41 | 1.36 | **1.68** |
| 10.0% $CO_2$, mTG | 0.58 | 1.11 | 2.56 | 0.35 | 0.44 | 0.93 | **0.52** |

**Fig. 1d and Supplementary Fig. 1l**

| | Median | | | FC (relative to Ambient) | | | **Geometric mean (FC)** |
|---|---|---|---|---|---|---|---|
| Condition | Rep_1 | Rep_2 | Rep_3 | Rep_1 | Rep_2 | Rep_3 | |
| Ambient, mTG | 0.26 | 0.18 | 0.22 | 1.00 | 1.00 | 1.00 | **1.00** |
| 2.0% $CO_2$, mTG | 3.15 | 2.72 | 4.27 | 12.03 | 15.41 | 19.62 | **15.38** |
| Ambient, Sugar-free | 0.05 | 0.05 | 0.06 | 1.00 | 1.00 | 1.00 | **1.00** |
| 2.0% $CO_2$, Sugar-free | 0.19 | 0.30 | 0.34 | 4.23 | 5.68 | 5.48 | **5.09** |
| Ambient, Nitrogen-free | 0.10 | 0.12 | 0.15 | 1.00 | 1.00 | 1.00 | **1.00** |
| 2.0% $CO_2$, Nitrogen-free | 0.44 | 1.06 | 2.94 | 4.30 | 8.49 | 19.35 | **8.90** |

| | Median | | | FC (relative to nutrient-free) | | | **Geometric mean (FC)** |
|---|---|---|---|---|---|---|---|
| Condition | Rep_1 | Rep_2 | Rep_3 | Rep_1 | Rep_2 | Rep_3 | |
| Ambient, Sugar-free | 0.05 | 0.05 | 0.06 | 1.00 | 1.00 | 1.00 | **1.00** |
| Ambient, mTG | 0.26 | 0.18 | 0.22 | 5.70 | 3.37 | 3.48 | **4.06** |
| 2.0% $CO_2$, Sugar-free | 0.19 | 0.30 | 0.34 | 1.00 | 1.00 | 1.00 | **1.00** |
| 2.0% $CO_2$, mTG | 3.15 | 2.72 | 4.27 | 16.22 | 9.16 | 12.46 | **12.28** |
| Ambient, Nitrogen-free | 0.10 | 0.12 | 0.15 | 1.00 | 1.00 | 1.00 | **1.00** |
| Ambient, mTG | 0.26 | 0.18 | 0.22 | 2.58 | 1.42 | 1.43 | **1.74** |
| 2.0% $CO_2$, Nitrogen-free | 0.44 | 1.06 | 2.94 | 1.00 | 1.00 | 1.00 | **1.00** |
| 2.0% $CO_2$, mTG | 3.15 | 2.72 | 4.27 | 7.23 | 2.57 | 1.45 | **3.00** |

### **Supplementary Table 2** (continued)

**Supplementary Fig. 1g**

| | Median | | | FC (relative to Ambient) | | | **Geometric mean (FC)** |
|---|---|---|---|---|---|---|---|
| Condition | Rep_1 | Rep_2 | Rep_3 | Rep_1 | Rep_2 | Rep_3 | |
| Ambient, Myr 500[†] | 0.78 | 0.95 | 0.60 | 1.00 | 1.00 | 1.00 | **1.00** |
| 2.0% $CO_2$, Myr 500[†] | 3.47 | 4.47 | 4.95 | 4.45 | 4.68 | 8.31 | **5.58** |
| Ambient, Myr 700 | 0.83 | 0.60 | 0.99 | 1.00 | 1.00 | 1.00 | **1.00** |
| 2.0% $CO_2$, Myr 700 | 4.56 | 5.40 | 6.49 | 5.53 | 8.99 | 6.59 | **6.89** |
| Ambient, Myr 1000 | 0.96 | 0.48 | 0.26 | 1.00 | 1.00 | 1.00 | **1.00** |
| 2.0% $CO_2$, Myr 1000 | 4.86 | 6.91 | 6.06 | 5.06 | 14.48 | 23.52 | **11.99** |
| Ambient, Pam 0 | 0.74 | 0.64 | 0.49 | 1.00 | 1.00 | 1.00 | **1.00** |
| 2.0% $CO_2$, Pam 0 | 3.36 | 3.80 | 4.73 | 4.53 | 5.91 | 9.68 | **6.38** |
| Ambient, Pam 200[†] | 0.78 | 0.95 | 0.60 | 1.00 | 1.00 | 1.00 | **1.00** |
| 2.0% $CO_2$, Pam 200[†] | 3.47 | 4.47 | 4.95 | 4.45 | 4.68 | 8.31 | **5.58** |
| Ambient, Pam 500 | 0.33 | 0.52 | 0.89 | 1.00 | 1.00 | 1.00 | **1.00** |
| 2.0% $CO_2$, Pam 500 | 2.21 | 3.64 | 4.76 | 6.78 | 6.94 | 5.36 | **6.32** |
| Ambient, PamΔ 0.1[†] | 0.78 | 0.95 | 0.60 | 1.00 | 1.00 | 1.00 | **1.00** |
| 2.0% $CO_2$, PamΔ 0.1[†] | 3.47 | 4.47 | 4.95 | 4.45 | 4.68 | 8.31 | **5.58** |
| Ambient, PamΔ 1 | 0.87 | 0.54 | 0.54 | 1.00 | 1.00 | 1.00 | **1.00** |
| 2.0% $CO_2$, PamΔ 1 | 3.60 | 4.06 | 4.81 | 4.12 | 7.49 | 8.97 | **6.52** |
| Ambient, PamΔ 10 | 0.11 | 0.08 | 0.15 | 1.00 | 1.00 | 1.00 | **1.00** |
| 2.0% $CO_2$, PamΔ 10 | 1.79 | 3.65 | 4.48 | 16.56 | 43.01 | 30.79 | **27.99** |

**Supplementary Fig. 1h**

| | Median | | | Fold change | | | **Geometric mean (FC)** |
|---|---|---|---|---|---|---|---|
| Condition | Rep_1 | Rep_2 | Rep_3 | Rep_1 | Rep_2 | Rep_3 | |
| Ambient, Myr 500[†] | 3.47 | 4.47 | 4.95 | 1.00 | 1.00 | 1.00 | **1.00** |
| Ambient, Myr 700 | 4.56 | 5.40 | 6.49 | 1.32 | 1.21 | 1.31 | **1.28** |
| Ambient, Myr 1000 | 4.86 | 6.91 | 6.06 | 1.40 | 1.55 | 1.22 | **1.38** |
| Ambient, Pam 0 | 3.36 | 3.80 | 4.73 | 1.00 | 1.00 | 1.00 | **1.00** |
| Ambient, Pam 200[†] | 3.47 | 4.47 | 4.95 | 1.03 | 1.18 | 1.05 | **1.08** |
| Ambient, Pam 500 | 2.21 | 3.64 | 4.76 | 0.66 | 0.96 | 1.01 | **0.86** |
| Ambient, PamΔ 0.1[†] | 3.47 | 4.47 | 4.95 | 1.00 | 1.00 | 1.00 | **1.00** |
| Ambient, PamΔ 1 | 3.60 | 4.06 | 4.81 | 1.04 | 0.91 | 0.97 | **0.97** |
| Ambient, PamΔ 10 | 1.79 | 3.65 | 4.48 | 0.52 | 0.82 | 0.91 | **0.73** |
| 2.0% $CO_2$, Myr 500[†] | 0.78 | 0.95 | 0.60 | 1.00 | 1.00 | 1.00 | **1.00** |
| 2.0% $CO_2$, Myr 700 | 0.83 | 0.60 | 0.99 | 1.06 | 0.63 | 1.65 | **1.03** |
| 2.0% $CO_2$, Myr 1000 | 0.96 | 0.48 | 0.26 | 1.23 | 0.50 | 0.43 | **0.64** |
| 2.0% $CO_2$, Pam 0 | 0.74 | 0.64 | 0.49 | 1.00 | 1.00 | 1.00 | **1.00** |
| 2.0% $CO_2$, Pam 200[†] | 0.78 | 0.95 | 0.60 | 1.05 | 1.48 | 1.22 | **1.24** |
| 2.0% $CO_2$, Pam 500 | 0.33 | 0.52 | 0.89 | 0.44 | 0.82 | 1.81 | **0.87** |
| 2.0% $CO_2$, PamΔ 0.1[†] | 0.78 | 0.95 | 0.60 | 1.00 | 1.00 | 1.00 | **1.00** |
| 2.0% $CO_2$, PamΔ 1 | 0.87 | 0.54 | 0.54 | 1.12 | 0.57 | 0.90 | **0.83** |
| 2.0% $CO_2$, PamΔ 10 | 0.11 | 0.08 | 0.15 | 0.14 | 0.09 | 0.24 | **0.14** |

**Supplementary Fig. 1n**

| | Median | | | Fold change | | | **Geometric mean (FC)** |
|---|---|---|---|---|---|---|---|
| Condition | Rep_1 | Rep_2 | Rep_3 | Rep_1 | Rep_2 | Rep_3 | |
| Ambient, mTGi | 0.62 | 0.96 | 0.99 | 1.00 | 1.00 | 1.00 | **1.00** |
| 0.5% $CO_2$, mTG | 0.62 | 1.12 | 0.77 | 1.01 | 1.16 | 0.78 | **0.97** |
| 2.0% $CO_2$, mTG | 0.34 | 0.32 | 0.66 | 0.55 | 0.33 | 0.66 | **0.49** |
| Ambient, pH 6.5 | 0.33 | 0.30 | 0.51 | 0.53 | 0.31 | 0.51 | **0.44** |
| Ambient, Myr 500 µM | 0.42 | 0.65 | 0.54 | 0.68 | 0.67 | 0.54 | **0.63** |
| Ambient, MES 10 mM | 0.45 | 0.49 | 0.45 | 0.73 | 0.51 | 0.45 | **0.55** |

**Supplementary Table 3.** Calculated values for statistical comparisons in this study.

**Fig. 1c, Ambient vs 2.0% $CO_2$ (Steel–Dwass test)**

| | *p*-value | | |
|---|---|---|---|
| Conditions | Rep_1 | Rep_2 | Rep_3 |
| 2.0% $CO_2$ -GR24 vs Ambient -GR24 | < 0.001 | < 0.001 | < 0.001 |
| 2.0% $CO_2$ -GR24 vs 0.5% $CO_2$ | 0.971 | < 0.001 | 1.000 |
| 2.0% $CO_2$ -GR24 vs 2.0% $CO_2$ | < 0.001 | < 0.001 | < 0.001 |
| 2.0% $CO_2$ -GR24 vs Ambient | 0.099 | 0.070 | < 0.001 |
| Ambient -GR24 vs 0.5% $CO_2$ | < 0.001 | < 0.001 | < 0.001 |
| Ambient -GR24 vs 2.0% $CO_2$ | < 0.001 | < 0.001 | < 0.001 |
| Ambient -GR24 vs Ambient | < 0.001 | < 0.001 | < 0.001 |
| 0.5% $CO_2$ vs 2.0% $CO_2$ | < 0.001 | < 0.001 | < 0.001 |
| 0.5% $CO_2$ vs Ambient | 0.001 | < 0.001 | < 0.001 |
| 2.0% $CO_2$ vs Ambient | < 0.001 | < 0.001 | < 0.001 |

**Fig. 1c, mTG vs GR24-free medium (Scheirer–Ray–Hare test)**

| | H statistic | | | *p*-value | | |
|---|---|---|---|---|---|---|
| Trials | 2.0% $CO_2$ | GR24 | $CO_2$ × GR24 | 2.0% $CO_2$ | GR24 | $CO_2$ × GR24 |
| Rep 1 | 38.39 | 14.96 | 0.01 | < 0.001 | < 0.001 | 0.918 |
| Rep 2 | 37.20 | 19.13 | 0.01 | < 0.001 | < 0.001 | 0.946 |
| Rep 3 | 50.15 | 14.10 | 0.06 | < 0.001 | < 0.001 | 0.810 |

**Fig. 1d, Ambient vs 2.0% $CO_2$ (Steel-Dwass test)**

| | *p*-value | | |
|---|---|---|---|
| Conditions | Rep_1 | Rep_2 | Rep_3 |
| Ambient mTG vs Ambient -Nitrogen | 0.054 | 0.500 | 1.000 |
| Ambient mTG vs Ambient -Sugar | < 0.001 | < 0.001 | 0.503 |
| Ambient mTG vs 2.0% $CO_2$ mTG | < 0.001 | < 0.001 | < 0.001 |
| Ambient mTG vs 2.0% $CO_2$ -Nitrogen | 0.962 | 0.057 | 0.001 |
| Ambient mTG vs 2.0% $CO_2$ -Sugar | 0.803 | 1.000 | 0.993 |
| Ambient -Nitrogen vs Ambient -Sugar | 0.003 | 0.180 | 0.078 |
| Ambient -Nitrogen vs 2.0% $CO_2$ mTG | < 0.001 | < 0.001 | < 0.001 |
| Ambient -Nitrogen vs 2.0% $CO_2$ -Nitrogen | 0.164 | < 0.001 | 0.004 |
| Ambient -Nitrogen vs 2.0% $CO_2$ -Sugar | 0.829 | 0.044 | 0.996 |
| Ambient -Sugar vs 2.0% $CO_2$ mTG | < 0.001 | < 0.001 | < 0.001 |
| Ambient -Sugar vs 2.0% $CO_2$ -Nitrogen | 0.023 | < 0.001 | < 0.001 |
| Ambient -Sugar vs 2.0% $CO_2$ -Sugar | 0.002 | < 0.001 | 0.002 |
| 2.0% $CO_2$ mTG vs 2.0% $CO_2$ -Nitrogen | < 0.001 | 0.036 | 0.004 |
| 2.0% $CO_2$ mTG vs 2.0% $CO_2$ -Sugar | < 0.001 | 0.017 | < 0.001 |
| 2.0% $CO_2$ -Nitrogen vs 2.0% $CO_2$ -Sugar | 0.296 | < 0.001 | 0.009 |

**Fig. 1d, mTG vs nutrient-free media (Scheirer-Ray-Hare test)**

| | H statistic | | | *p*-value | | |
|---|---|---|---|---|---|---|
| Trials | 2.0% $CO_2$ | Nutrient | $CO_2$ × nutrient | 2.0% $CO_2$ | Nutrient | $CO_2$ × nutrient |
| Rep_1 | 18.74 | 27.16 | 0.42 | < 0.001 | < 0.001 | 0.809 |
| Rep_2 | 30.64 | 14.38 | 0.24 | < 0.001 | < 0.001 | 0.888 |
| Rep_3 | 33.85 | 17.23 | 3.73 | < 0.001 | < 0.001 | 0.154 |

### **Supplementary Table 3** (continued)

**Supplementary Fig. 1d, 5.0 or10.0% $CO_2$ vs 2.0% $CO_2$ (Steel-Dwass test)**

| | *p*-value | | |
|---|---|---|---|
| Conditions | Rep_1 | Rep_2 | Rep_3 |
| 10.0% $CO_2$ vs 2.0% $CO_2$ | 0.014 | 0.040 | 0.588 |
| 10.0% $CO_2$ vs 5.0% $CO_2$ | < 0.001 | 0.001 | < 0.001 |
| 2.0% $CO_2$ vs 5.0% $CO_2$ | 0.126 | 0.365 | 0.134 |

**Supplementary Fig. 1e, Ambient vs 2.0% $CO_2$ (Wilcoxon rank-sum test)**

| | *p*-values | | |
|---|---|---|---|
| | Rep_1 | Rep_2 | Rep_3 |
| Maximum Feret diameter | < 0.001 | 0.835 | 0.497 |
| Minimum Feret diameter | 0.207 | 0.403 | 0.183 |

**Supplementary Fig 1f, The lowest vs higher concentration of each fatty acid (Steel test)**

| | *p*-values | | | | | |
|---|---|---|---|---|---|---|
| | Ambient | | | 2.0% $CO_2$ | | |
| | Rep_1 | Rep_2 | Rep_3 | Rep_1 | Rep_2 | Rep_3 |
| Myr 500 µM vs Myr 700 µM | 0.852 | 0.212 | 0.462 | < 0.001 | 0.619 | < 0.001 |
| Myr 500 µM vs Myr 1000 µM | 0.969 | 0.085 | 0.030 | < 0.001 | 0.001 | 0.083 |
| Pam 0 µM vs Pam 200 µM | 0.959 | 0.524 | 0.736 | 0.994 | 0.056 | 0.548 |
| Pam 0 µM vs Pam 500 µM | 0.024 | 0.219 | 0.558 | 0.002 | 0.976 | 0.987 |
| PamΔ 0.1 µM vs PamΔ 1 µM | 0.983 | 0.165 | 0.998 | 0.640 | 0.194 | 0.436 |
| PamΔ 0.1 µM vs PamΔ 10 µM | < 0.001 | < 0.001 | 0.003 | 0.004 | 0.090 | 0.031 |

**Fig. 1e, mTG under ambient control vs various conditions (Steel test)**

| | *p*-values | | |
|---|---|---|---|
| | Rep_1 | Rep_2 | Rep_3 |
| 0.5% $CO_2$ | 1.000 | 0.829 | 0.331 |
| 2.0% $CO_2$ | < 0.001 | < 0.001 | < 0.001 |
| pH 6.5 | < 0.001 | < 0.001 | < 0.001 |
| Myr 500 µM | 0.007 | < 0.001 | < 0.001 |
| MES 10 mM | 0.012 | < 0.001 | < 0.001 |

**Supplementary Table 4.** Summary of parent spore viability in culture experiments

| Experiment | Trial | Atmospheric condition | Medium | All incubated spores | Contaminated | Ungerminated | Sample size |
|---|---|---|---|---|---|---|---|
| *R. clarus* in media with various concentration of fatty acids (Supplementary Fig. 1f–h) | 1 | Ambient | Pam_0 | 20 | 0 | 0 | 20 |
| | | | Pam_200 (=mTG) | 20 | 0 | 0 | 20 |
| | | | Pam_500 | 20 | 0 | 0 | 20 |
| | | | PamΔ_1 | 20 | 0 | 0 | 20 |
| | | | PamΔ_10 | 20 | 0 | 0 | 20 |
| | | | Myr_700 | 20 | 0 | 0 | 20 |
| | | | Myr_1000 | 20 | 0 | 0 | 20 |
| | | $CO_2$ 2.0% | Pam_0 | 20 | 0 | 0 | 20 |
| | | | Pam_200 (mTG) | 20 | 0 | 0 | 20 |
| | | | Pam_500 | 20 | 0 | 0 | 20 |
| | | | PamΔ_1 | 20 | 0 | 0 | 20 |
| | | | PamΔ_10 | 20 | 0 | 1 | 19 |
| | | | Myr_700 | 20 | 0 | 0 | 20 |
| | | | Myr_1000 | 20 | 0 | 0 | 20 |
| | 2 | Ambient | Pam_0 | 16 | 0 | 0 | 16 |
| | | | Pam_200 (mTG) | 16 | 0 | 0 | 16 |
| | | | Pam_500 | 16 | 0 | 0 | 16 |
| | | | PamΔ_1 | 16 | 0 | 0 | 16 |
| | | | PamΔ_10 | 16 | 0 | 0 | 16 |
| | | | Myr_700 | 16 | 0 | 0 | 16 |
| | | | Myr_1000 | 16 | 0 | 0 | 16 |
| | | $CO_2$ 2.0% | Pam_0 | 16 | 0 | 0 | 16 |
| | | | Pam_200 (mTG) | 16 | 0 | 0 | 16 |
| | | | Pam_500 | 16 | 0 | 0 | 16 |
| | | | PamΔ_1 | 16 | 0 | 0 | 16 |
| | | | PamΔ_10 | 16 | 1 | 0 | 15 |
| | | | Myr_700 | 16 | 0 | 1 | 15 |
| | | | Myr_1000 | 16 | 0 | 0 | 16 |
| | 3 | Ambient | Pam_0 | 15 | 0 | 0 | 15 |
| | | | Pam_200 (mTG) | 15 | 0 | 0 | 15 |
| | | | Pam_500 | 15 | 0 | 0 | 15 |
| | | | PamΔ_1 | 15 | 0 | 0 | 15 |
| | | | PamΔ_10 | 15 | 0 | 0 | 15 |
| | | | Myr_700 | 15 | 0 | 0 | 15 |
| | | | Myr_1000 | 15 | 0 | 0 | 15 |
| | | $CO_2$ 2.0% | Pam_0 | 15 | 0 | 0 | 15 |
| | | | Pam_200 (mTG) | 15 | 0 | 0 | 15 |
| | | | Pam_500 | 15 | 0 | 0 | 15 |
| | | | PamΔ_1 | 15 | 0 | 0 | 15 |
| | | | PamΔ_10 | 15 | 0 | 0 | 15 |
| | | | Myr_700 | 15 | 0 | 0 | 15 |
| | | | Myr_1000 | 15 | 1 | 0 | 14 |
| *R. clarus* in the mTG and GR24-free media (Fig. 1c and Supplementary Fig. 1b) | 1 | Ambient | mTG - GR24 | 20 | 0 | 0 | 20 |
| | | $CO_2$ 2.0% | | 20 | 0 | 0 | 20 |
| | | Ambient | mTG | 20 | 0 | 0 | 20 |
| | | $CO_2$ 0.5% | | 20 | 1 | 0 | 19 |
| | | $CO_2$ 2.0% | | 20 | 0 | 0 | 20 |
| | 2 | Ambient | mTG - GR24 | 20 | 1 | 1 | 18 |
| | | $CO_2$ 2.0% | | 20 | 1 | 0 | 19 |
| | | Ambient | mTG | 20 | 1 | 0 | 19 |
| | | $CO_2$ 0.5% | | 20 | 0 | 0 | 20 |
| | | $CO_2$ 2.0% | | 20 | 1 | 0 | 19 |
| | 3 | Ambient | mTG - GR24 | 20 | 0 | 0 | 20 |
| | | $CO_2$ 2.0% | | 20 | 0 | 1 | 19 |
| | | Ambient | mTG | 20 | 1 | 1 | 18 |
| | | $CO_2$ 0.5% | | 20 | 0 | 0 | 20 |
| | | $CO_2$ 2.0% | | 20 | 0 | 0 | 20 |

**Supplementary Table 4** (continued)

| Experiment | Trial | Atmospheric condition | Medium | All incubated spores | Contaminated | Ungerminated or quite bad growth | Sample size |
|---|---|---|---|---|---|---|---|
| *R. clarus* under high $CO_2$ conditions (Supplementary Fig. 1d) | 1 | $CO_2$ 2.0% | mTG | 16 | 4 | 0 | 12 |
| | | $CO_2$ 5.0% | | 16 | 3 | 0 | 13 |
| | | $CO_2$ 10.0% | | 16 | 1 | 0 | 15 |
| | 2 | $CO_2$ 2.0% | | 16 | 0 | 0 | 16 |
| | | $CO_2$ 5.0% | | 16 | 0 | 1 | 15 |
| | | $CO_2$ 10.0% | | 17 | 0 | 2 | 15 |
| | 3 | $CO_2$ 2.0% | | 16 | 0 | 0 | 16 |
| | | $CO_2$ 5.0% | | 16 | 0 | 0 | 16 |
| | | $CO_2$ 10.0% | | 17 | 0 | 0 | 17 |
| *R. clarus* in fatty acid-free media (Supplementary Fig. 1e) | 1 | Ambient | mTG - fatty acids | 12 | 0 | 1 | 11 |
| | | $CO_2$ 2.0% | | 12 | 0 | 0 | 12 |
| | 2 | Ambient | | 12 | 7 | 0 | 5 |
| | | $CO_2$ 2.0% | | 12 | 7 | 0 | 5 |
| | 3 | Ambient | | 13 | 0 | 0 | 13 |
| | | $CO_2$ 2.0% | | 12 | 0 | 0 | 12 |
| *R. clarus* in sugar- or nitrogen-free media (Fig. 1d, Supplementary Fig. 1l) | 1 | Ambient | mTG | 12 | 0 | 0 | 12 |
| | | | mTG - sugars | 12 | 0 | 1 | 11 |
| | | | mTG - nitrogen | 12 | 0 | 1 | 11 |
| | | $CO_2$ 2.0% | mTG | 12 | 0 | 1 | 11 |
| | | | mTG - sugars | 12 | 0 | 1 | 11 |
| | | | mTG - nitrogen | 12 | 0 | 0 | 12 |
| | 2 | Ambient | mTG | 25 | 16 | 0 | 9 |
| | | | mTG - sugars | 25 | 17 | 0 | 8 |
| | | | mTG - nitrogen | 25 | 14 | 0 | 11 |
| | | $CO_2$ 2.0% | mTG | 25 | 14 | 1 | 10 |
| | | | mTG - sugars | 25 | 16 | 0 | 9 |
| | | | mTG - nitrogen | 25 | 13 | 0 | 12 |
| | 3 | Ambient | mTG | 15 | 0 | 0 | 15 |
| | | | mTG - sugars | 15 | 0 | 0 | 15 |
| | | | mTG - nitrogen | 15 | 0 | 0 | 15 |
| | | $CO_2$ 2.0% | mTG | 15 | 0 | 0 | 15 |
| | | | mTG - sugars | 15 | 0 | 0 | 15 |
| | | | mTG - nitrogen | 15 | 0 | 0 | 15 |
| *R. irregularis* in various conditions (Fig. 1e, Supplementary Fig. 1n) | 1 | Ambient | mTGi | 15 | 0 | 0 | 15 |
| | | $CO_2$ 0.5% | | 15 | 0 | 0 | 15 |
| | | $CO_2$ 2.0% | | 15 | 0 | 0 | 15 |
| | | Ambient | mTGi Myr 500 µM | 15 | 0 | 0 | 15 |
| | | | mTGi pH 6.5 | 15 | 0 | 0 | 15 |
| | | | mTGi MES 10 mM | 15 | 0 | 0 | 15 |
| | 2 | Ambient | mTGi | 18 | 0 | 0 | 18 |
| | | $CO_2$ 0.5% | | 18 | 0 | 0 | 18 |
| | | $CO_2$ 2.0% | | 18 | 0 | 0 | 18 |
| | | Ambient | mTGi Myr 500 µM | 15 | 0 | 0 | 15 |
| | | | mTGi pH 6.5 | 14 | 0 | 0 | 14 |
| | | | mTGi MES 10 mM | 15 | 0 | 0 | 15 |
| | 3 | Ambient | mTGi | 15 | 0 | 0 | 15 |
| | | $CO_2$ 0.5% | | 15 | 0 | 0 | 15 |
| | | $CO_2$ 2.0% | | 15 | 0 | 0 | 15 |
| | | Ambient | mTGi Myr 500 µM | 15 | 0 | 0 | 15 |
| | | | mTGi pH 6.5 | 15 | 0 | 0 | 15 |
| | | | mTGi MES 10 mM | 15 | 0 | 0 | 15 |

a

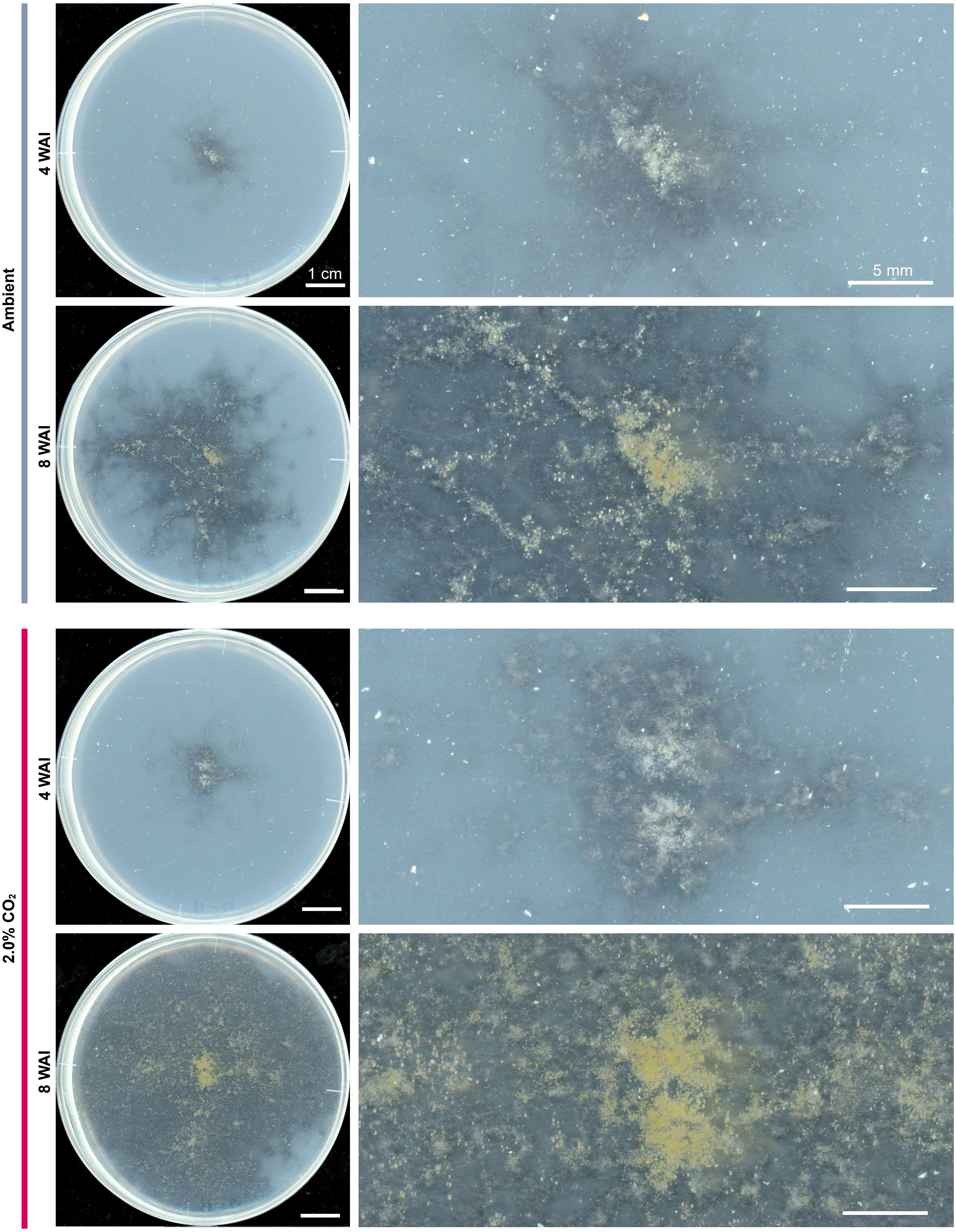

**b**

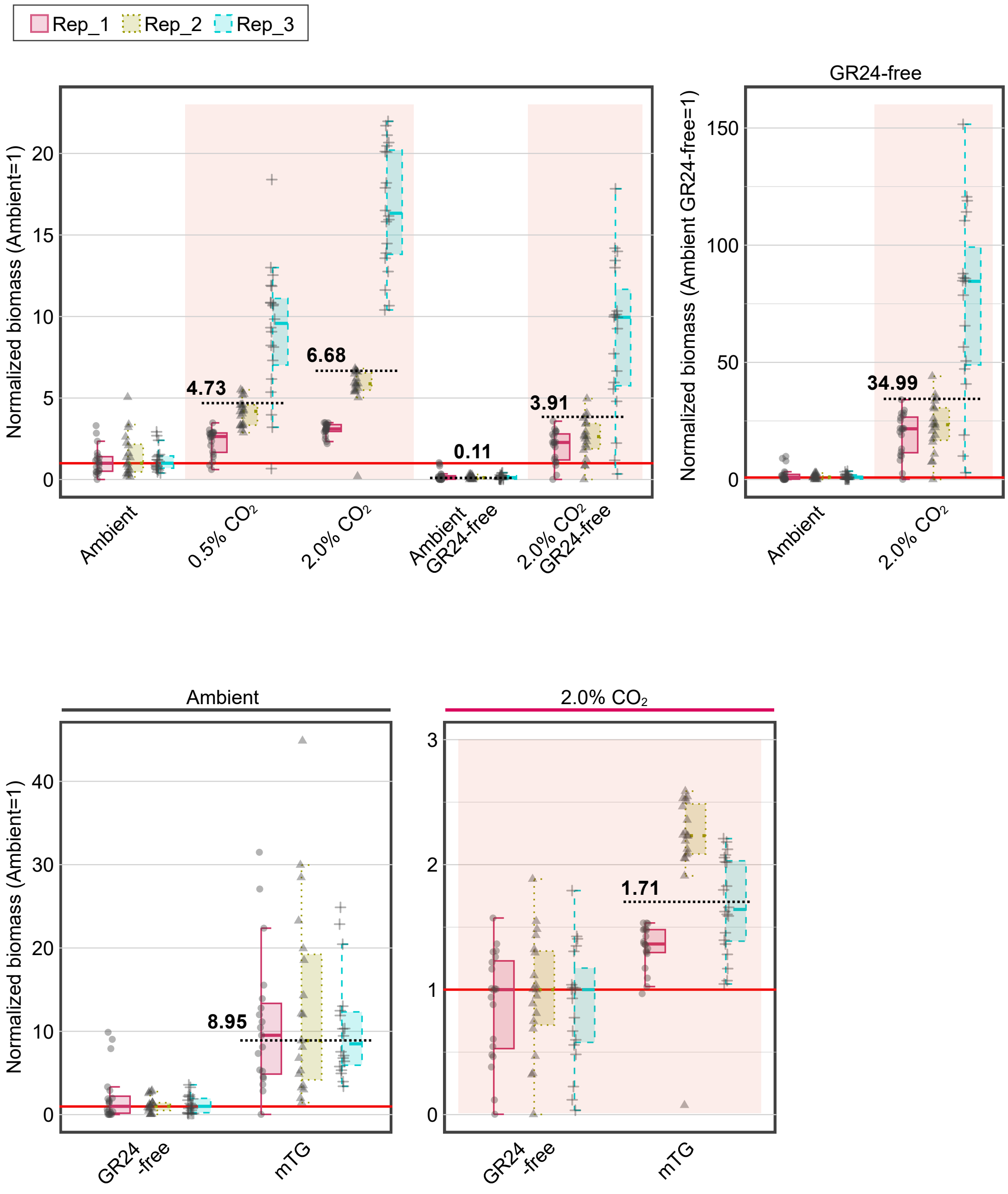
Rep_1
Rep_2
Rep_3
Normalized biomass (Ambient=1)
4.73
6.68
0.11
3.91
Ambient
0.5% CO2
2.0% CO2
Ambient GR24-free
2.0% CO2 GR24-free
GR24-free
Normalized biomass (Ambient GR24-free=1)
34.99
Ambient
2.0% CO2
Ambient
2.0% CO2
8.95
1.71
GR24 -free
mTG

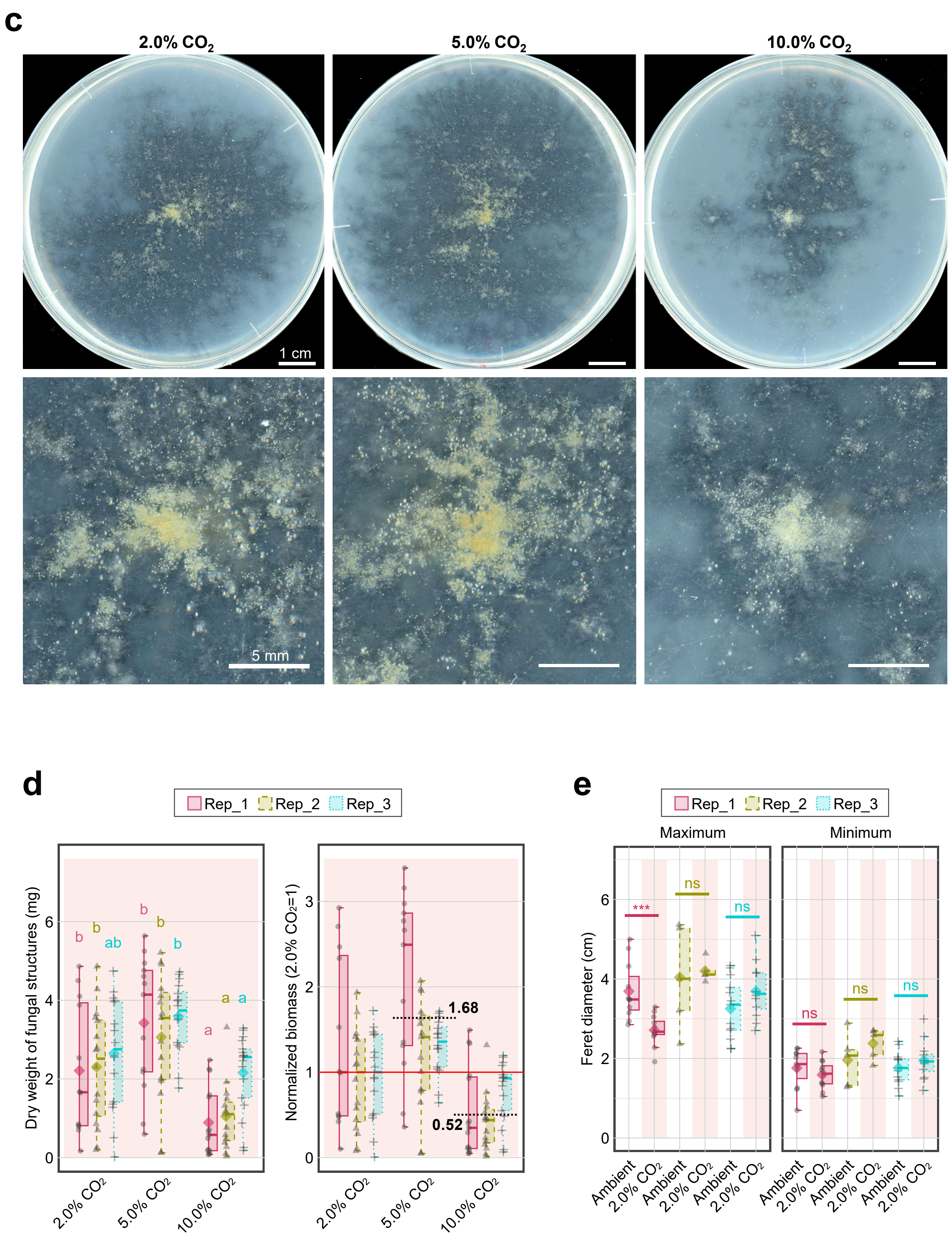

c
2.0% CO2
5.0% CO2
10.0% CO2
1 cm
5 mm
d
Rep_1
Rep_2
Rep_3
Dry weight of fungal structures (mg)
Normalized biomass (2.0% CO2=1)
1.68
0.52
2.0% CO2
5.0% CO2
10.0% CO2
e
Maximum
Minimum
Feret diameter (cm)
***
ns
Ambient

f

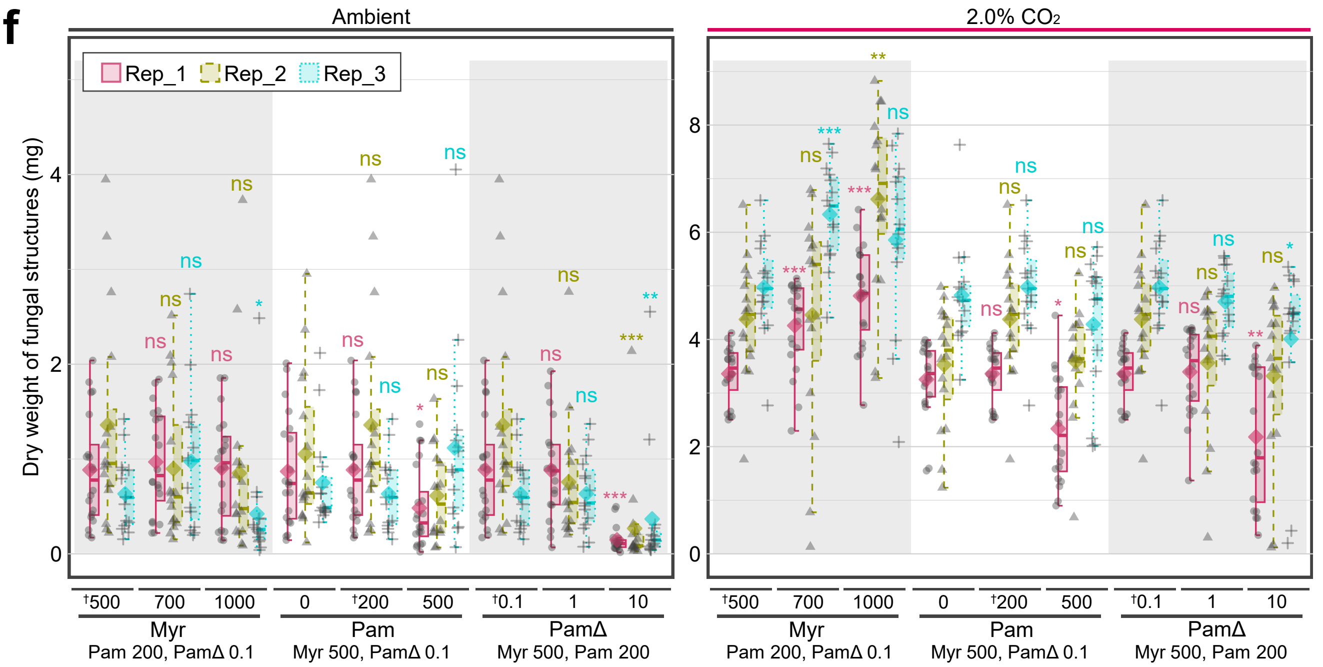

Ambient
2.0% CO2
Rep_1 Rep_2 Rep_3
Dry weight of fungal structures (mg)
†500 700 1000 0 †200 500 †0.1 1 10
Myr
Pam 200, PamΔ 0.1
Pam
Myr 500, PamΔ 0.1
PamΔ
Myr 500, Pam 200


g

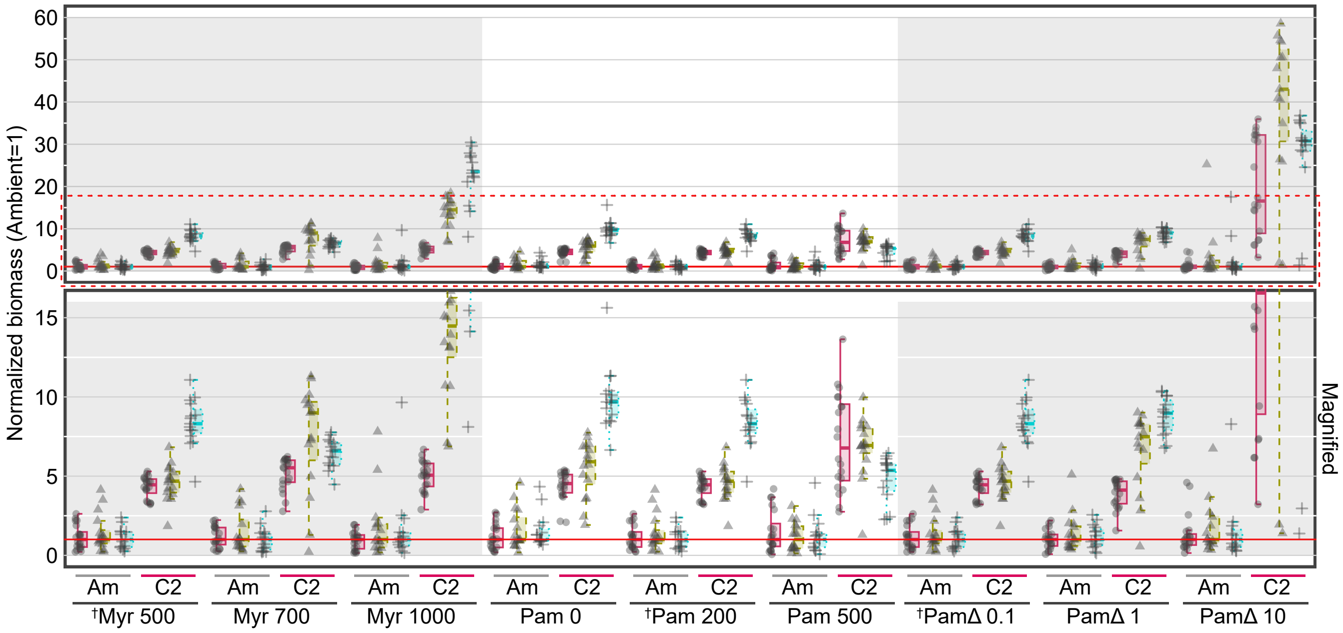

Normalized biomass (Ambient=1)
Magnified
Am C2
†Myr 500
Myr 700
Myr 1000
Pam 0
†Pam 200
Pam 500
†PamΔ 0.1
PamΔ 1
PamΔ 10


h

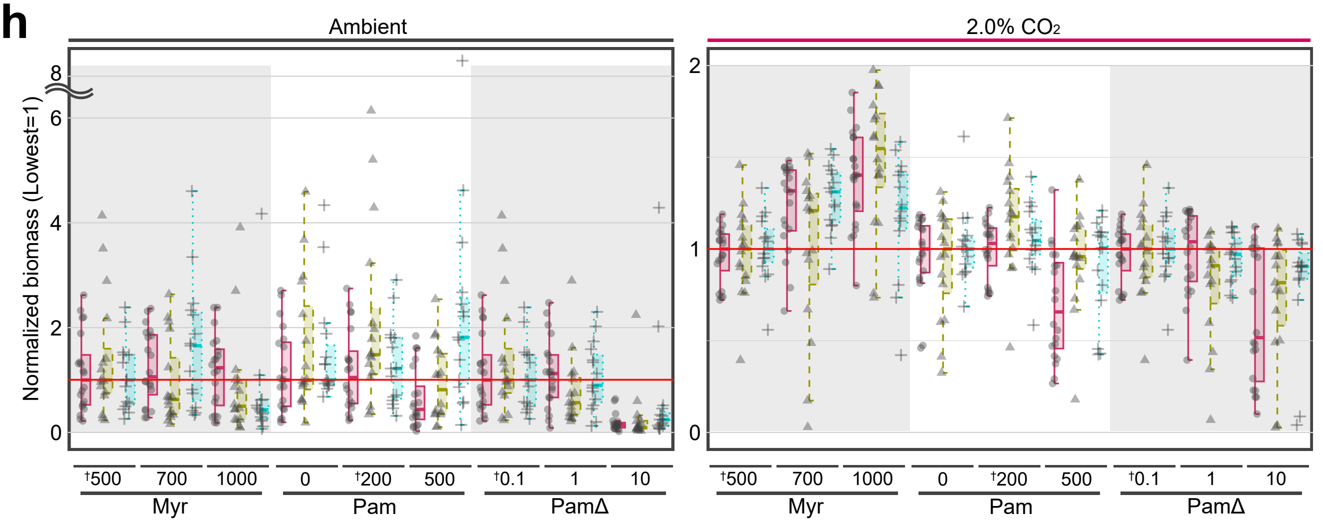

Ambient
2.0% CO2
Normalized biomass (Lowest=1)
†500 700 1000 0 †200 500 †0.1 1 10
Myr
Pam
PamΔ

i

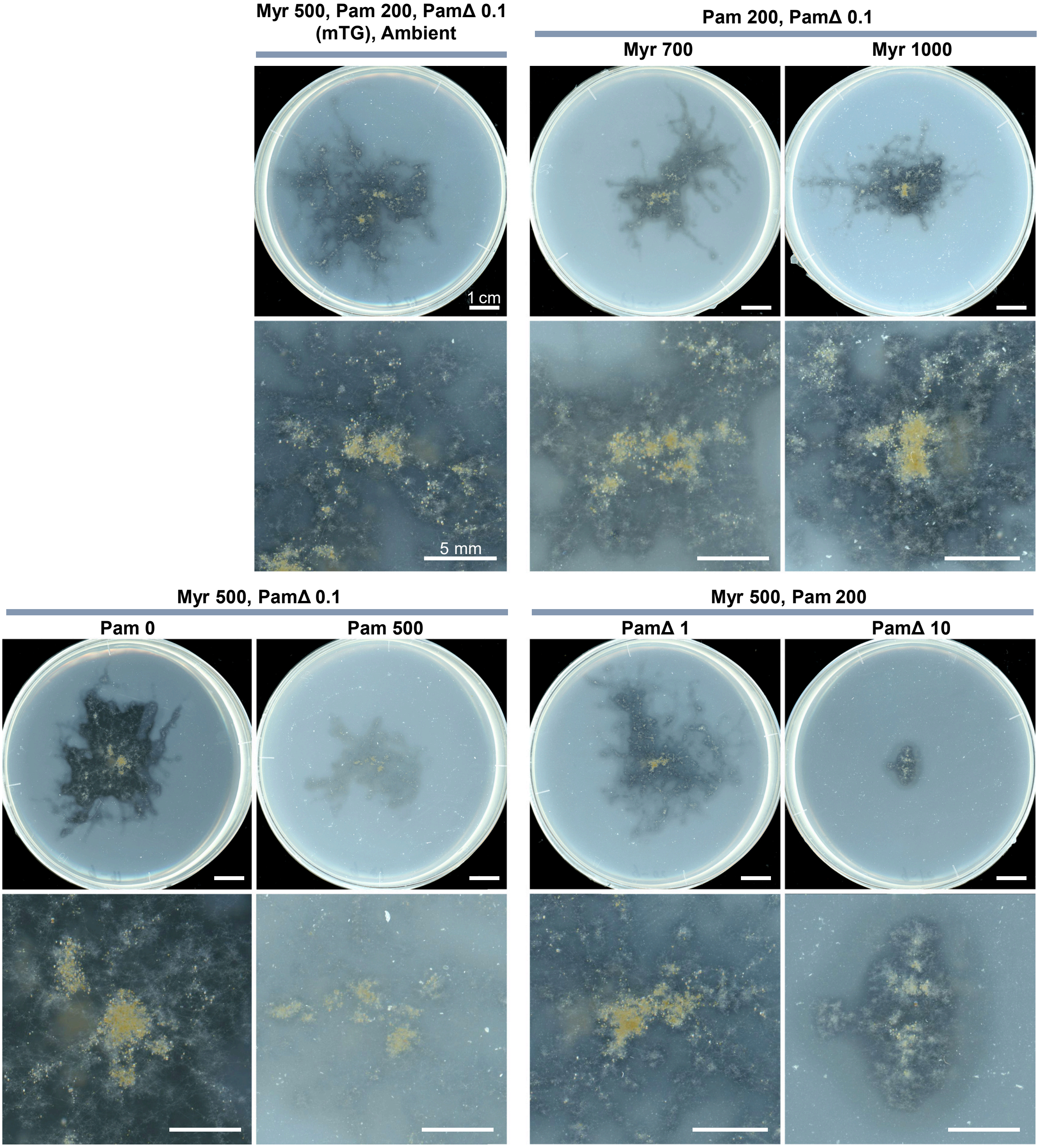

Myr 500, Pam 200, PamΔ 0.1
(mTG), Ambient
Pam 200, PamΔ 0.1
Myr 700
Myr 1000
1 cm
5 mm
Myr 500, PamΔ 0.1
Pam 0
Pam 500
Myr 500, Pam 200
PamΔ 1
PamΔ 10

j

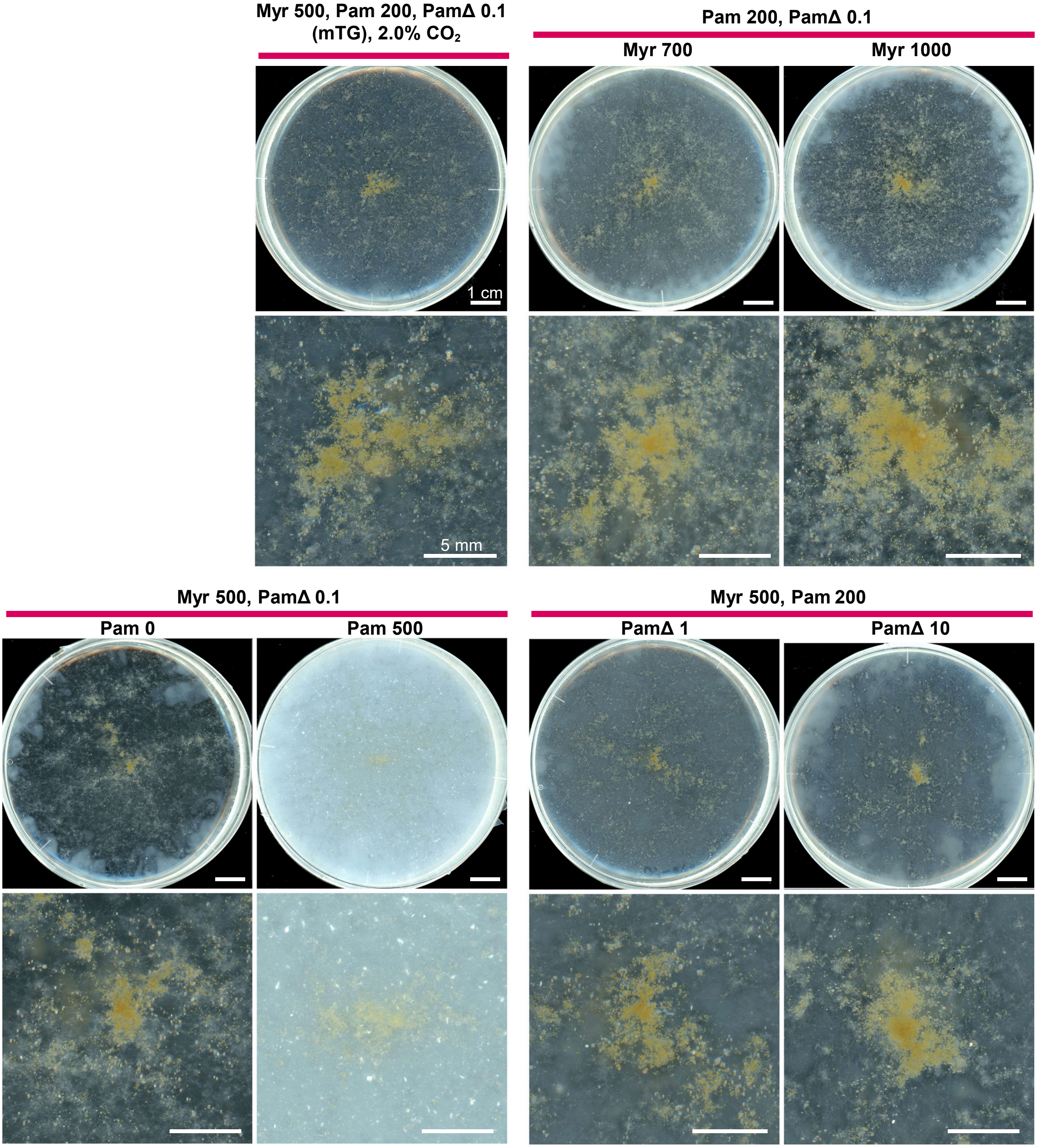

Myr 500, Pam 200, PamΔ 0.1
(mTG), 2.0% CO2
Pam 200, PamΔ 0.1
Myr 700
Myr 1000
1 cm
5 mm
Myr 500, PamΔ 0.1
Pam 0
Pam 500
Myr 500, Pam 200
PamΔ 1
PamΔ 10

k

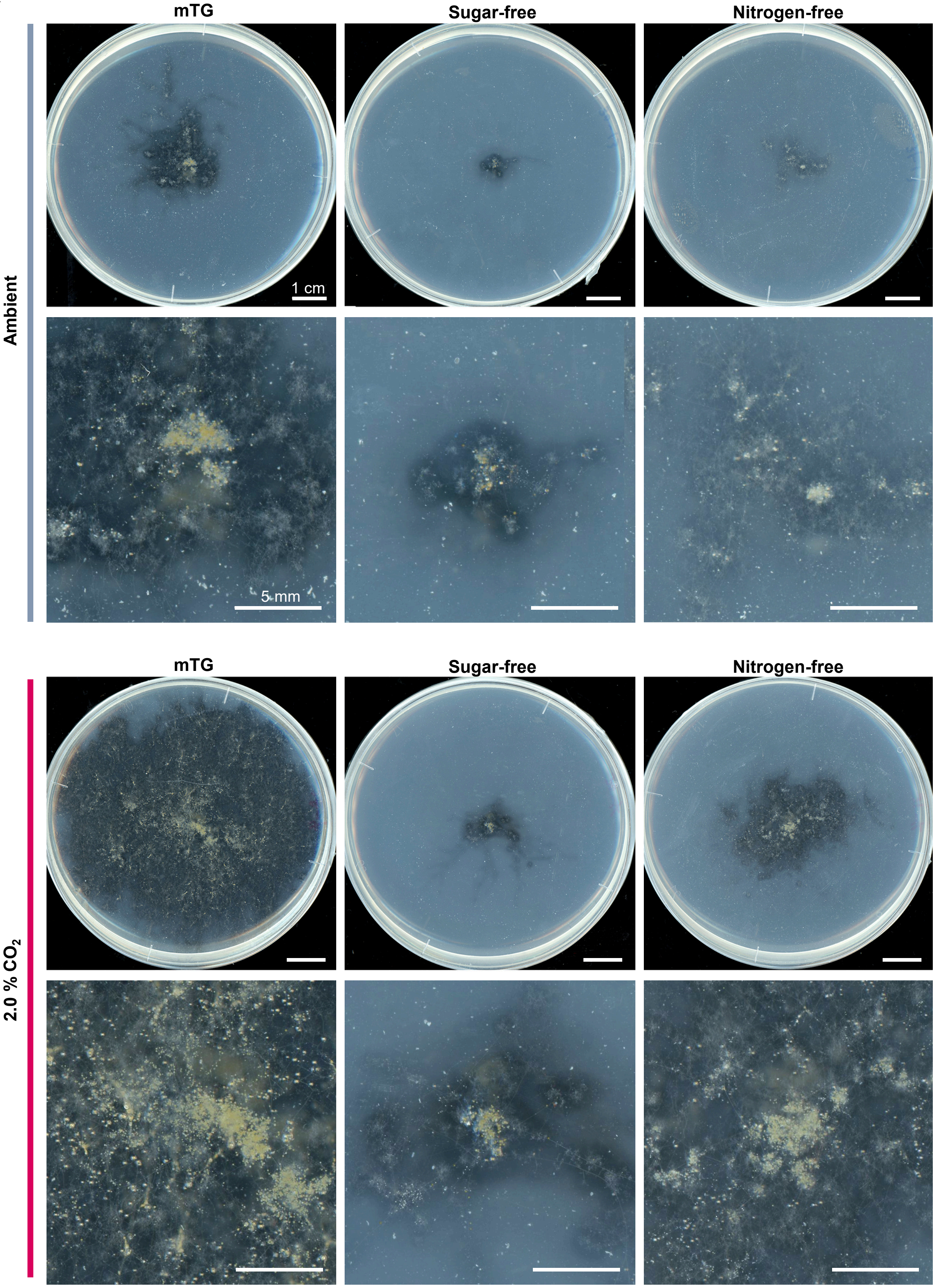

mTG
Sugar-free
Nitrogen-free
Ambient
1 cm
5 mm
mTG
Sugar-free
Nitrogen-free
2.0 % CO2

**I**

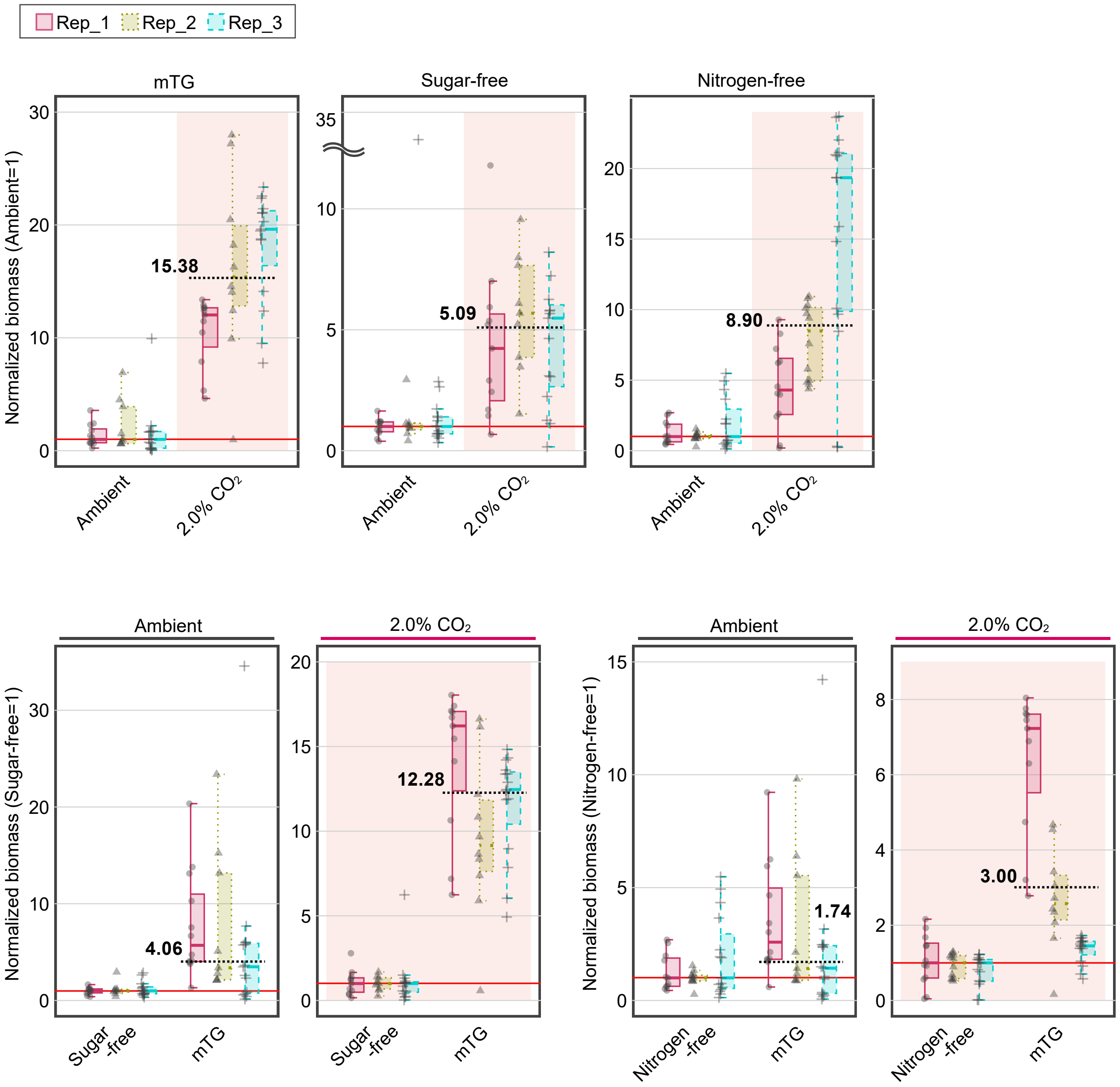

Rep_1
Rep_2
Rep_3
mTG
Sugar-free
Nitrogen-free
Normalized biomass (Ambient=1)
15.38
5.09
8.90
Ambient
2.0% CO2
Ambient
2.0% CO2
Ambient
2.0% CO2
Normalized biomass (Sugar-free=1)
4.06
12.28
Sugar -free
mTG
Normalized biomass (Nitrogen-free=1)
1.74
3.00
Nitrogen -free
mTG

**m**

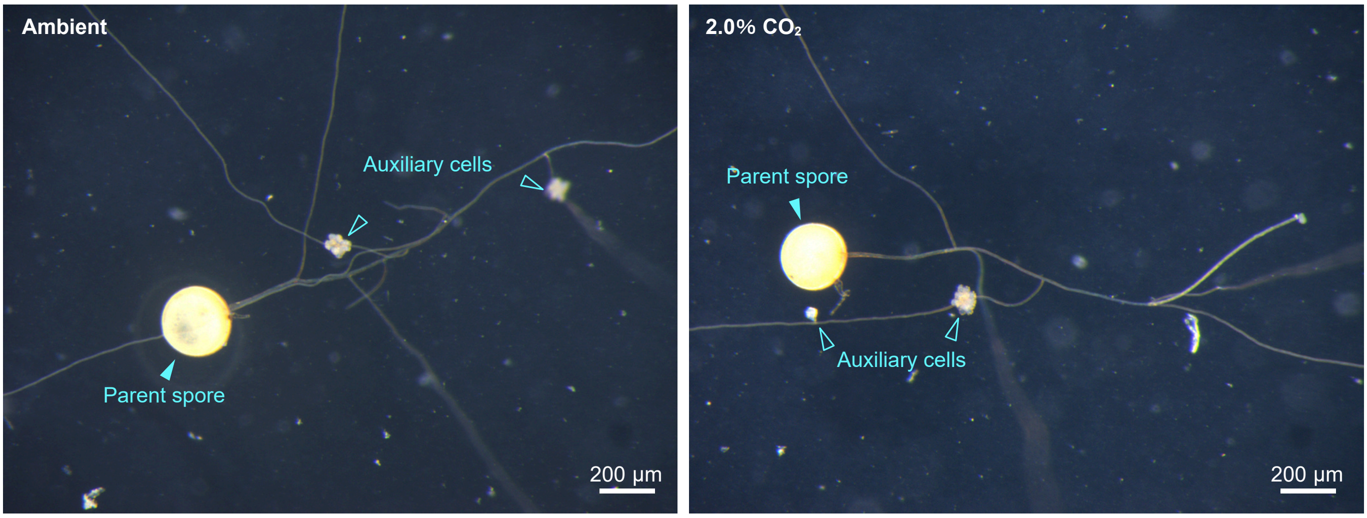

Ambient
Auxiliary cells
Parent spore
200 µm
2.0% CO2
Parent spore
Auxiliary cells
200 µm


**n**

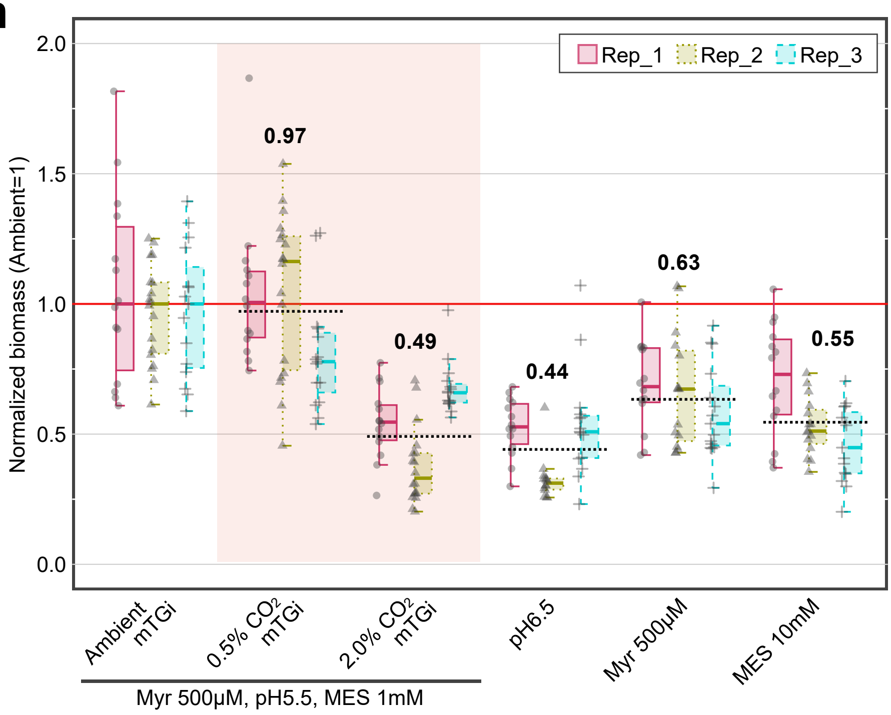

Rep_1
Rep_2
Rep_3
Normalized biomass (Ambient=1)
2.0
1.5
1.0
0.5
0.0
0.97
0.49
0.44
0.63
0.55
Ambient mTGi
0.5% CO2 mTGi
2.0% CO2 mTGi
pH6.5
Myr 500µM
MES 10mM
Myr 500µM, pH5.5, MES 1mM

**o**

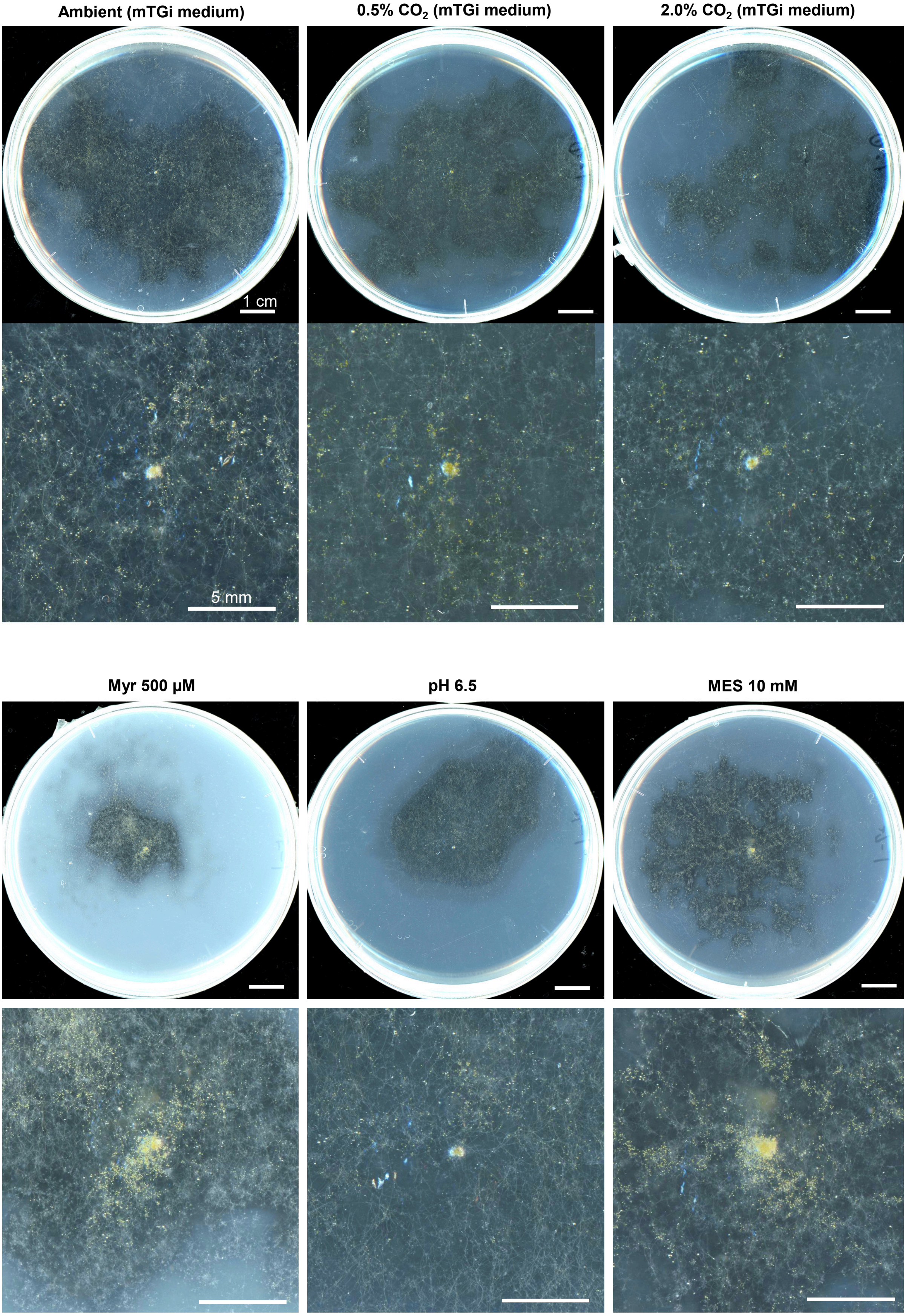

Ambient (mTGi medium)
0.5% $CO_2$ (mTGi medium)
2.0% $CO_2$ (mTGi medium)
1 cm
5 mm
Myr 500 µM
pH 6.5
MES 10 mM

**p**

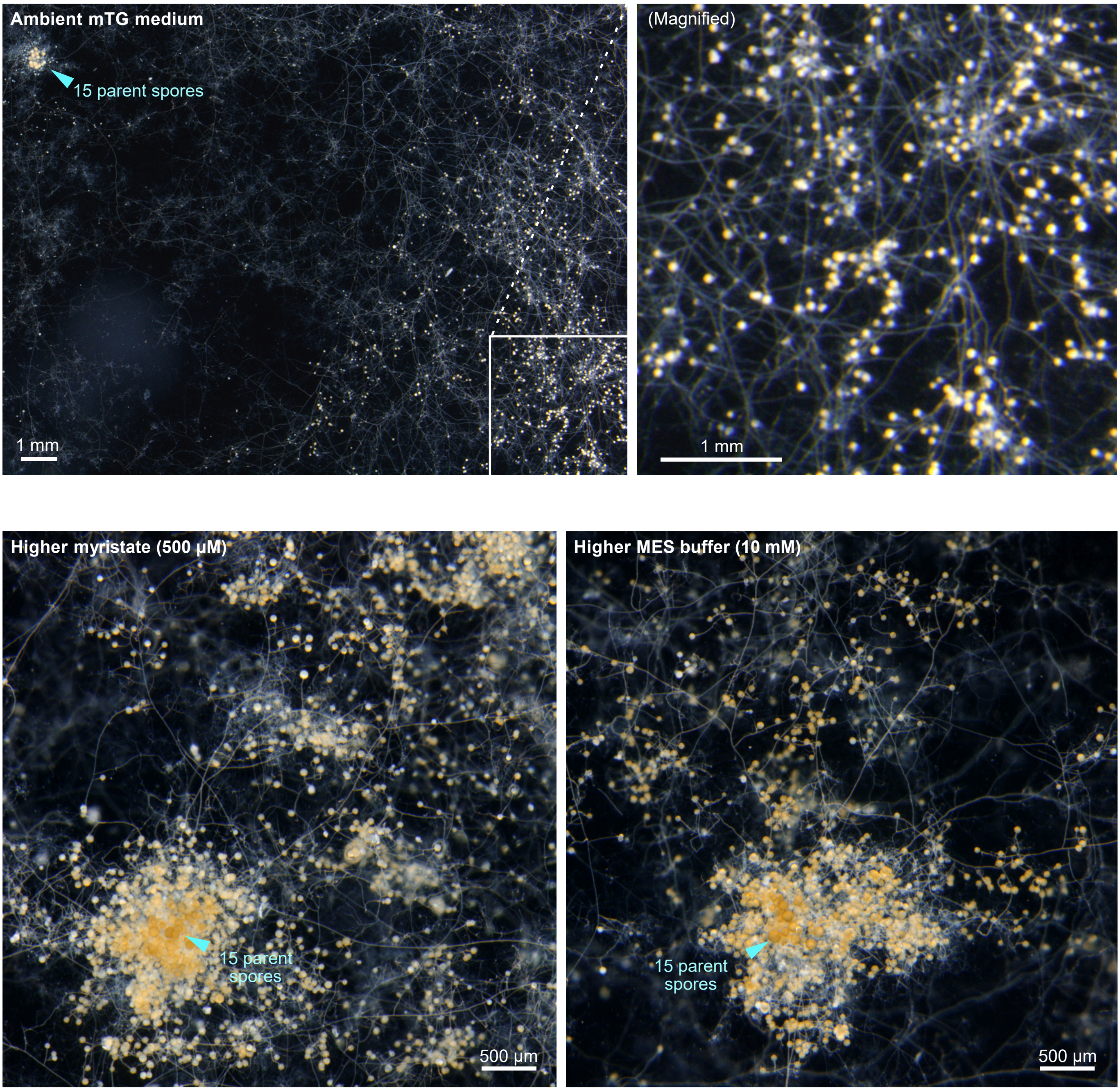

Ambient mTG medium
15 parent spores
1 mm
(Magnified)
1 mm
Higher myristate (500 µM)
15 parent spores
500 µm
Higher MES buffer (10 mM)
15 parent spores
500 µm

**Supplementary Figure 1. Asymbiotic growth of AM fungi under various conditions.**

**a,** Representative images of asymbiotically growing *R. clarus* in mTG medium at 4 and 8 WAI. Top panels show whole dishes, and bottom panels show magnified images around parent spores. Bars, 1 cm (top) and 5 mm (bottom). **b**, Relative biomass fold-changes, normalised to the median value of the ambient mTG or ambient GR24-free conditions (top panels), and to the median value of GR24-free controls under each atmospheric condition (bottom panels), for each medium composition. Red lines indicate the normalised control level (= 1), and black dotted lines indicate geometric means, with the corresponding values shown alongside. Median values, fold-changes and geometric means are provided in Supplementary Table 2. **c**–**d**, Effects of higher $CO_2$ concentrations on *R. clarus*. Representative images of asymbiotically growing *R. clarus* under 2.0, 5.0, and 10.0% $CO_2$ (**c**). Top panels show whole dishes, and bottom panels show magnified images around parent spores. Bars, 1 cm (top) and 5 mm (bottom). Dry weight of fungal structures (**d**, left) and relative biomass fold-changes (**d**, right) under 2.0, 5.0, and 10.0% $CO_2$. Diamonds indicate means. Different letters indicate significant differences among conditions. Statistical significance was determined using the Steel–Dwass test ($p < 0.05$). A red line indicates the normalised control level (= 1), and black dotted lines indicate geometric means, with the corresponding values shown alongside. Median values, fold-changes and geometric means are provided in Supplementary Table 2. **e**, Comparison of colony size in asymbiotically cultured *R. clarus* under ambient and 2.0% $CO_2$ conditions in fatty acid-free medium using Feret diameters. Diamonds indicate means. Statistical significance was determined using the Wilcoxon rank-sum test ($p < 0.05$). ***, $p < 0.001$; ns, not significant. **f**–**j**, Contribution of fatty acids and 2.0% $CO_2$ on *R. clarus* growth. In each fatty acid series, the concentrations of two fatty acids were fixed to evaluate the effects of the focused fatty acid. Daggers (†) represent the same sample (mTG medium) used as the common reference for each series. Myr, potassium myristate; Pam, potassium palmitate; PamΔ, potassium palmitoleate. Numbers following fatty acids indicate their concentrations in µM. Dry weight of fungal structures under ambient and 2.0% $CO_2$ conditions across various combinations of fatty acids (**f**). Diamonds indicate means. Statistical significance was determined using the Steel test ($p < 0.05$) within each series, with the lowest concentration serving as the control. ***, $p < 0.001$; **, $0.001 \leq p < 0.01$ ***; $0.01 \leq p < 0.05$; ns, not significant. Relative biomass fold-changes (**g**–**h**), normalised to the median value of the ambient condition for each medium composition (**g**) and to the median value of the lowest concentration for each fatty acid series (**h**). The lower panel in **g** shows an enlarged view. Am, Ambient; C2, 2.0% $CO_2$. Red lines indicate the normalised control level (= 1). Median values and fold-changes are also provided in Supplementary Table 2. Representative images of asymbiotically growing *R. clarus* in media with various concentrations of fatty acids under ambient (**i**) and 2.0% $CO_2$ (**j**) conditions. Top panels show whole dishes, and bottom panels show magnified images around parent spores. Bars, 1 cm (top) and 5 mm (bottom). **k**, **l**, Effects of sugars and nitrogen under 2.0% $CO_2$ on *R. clarus* growth. Representative images of asymbiotically growing *R. clarus* in the mTG and sugar- or nitrogen-free media (**k**). Top panels show whole dishes, and bottom panels show magnified images around parent spores. Bars, 1 cm (top) and 5 mm (bottom). Relative biomass fold-changes (**l**), normalised to the median value of the ambient condition for each medium (top panels), and to the median value of each nutrient-free medium control under each atmospheric condition (bottom panles). Red lines indicate the normalised control level (= 1), and black dotted lines indicate geometric means, with the corresponding values shown alongside. Median values, fold-changes and geometric means are provided in Supplementary Table 2. **m**, Asymbiotic growing *G.rosea* forming auxiliary cells in the presence of fatty acids under ambient and 2.0% $CO_2$ conditions. **n**–**p**, Asymbiotic growth of *R. irregularis*. Relative biomass fold-changes normalised to the median value of the ambient mTGi control for each condition (**n**). A red line indicates the normalised control level (= 1), and black dotted lines indicate geometric means, with the corresponding values shown alongside. Median values, fold-changes and geometric means are provided in Supplementary Table 2. **o**, Representative images of asymbiotically growing *R. irregularis* in various conditions. Top panels show whole dishes, and bottom panels show magnified images around parent spores. Bars, 1 cm (top) and 5 mm (bottom). **p**, Morphological patterns frequently observed in *R. irregularis*. Spore formation at a distance from parent spores (top), and spore production around parent spores in the medium with higher concentrations of myristate or MES (bottom).

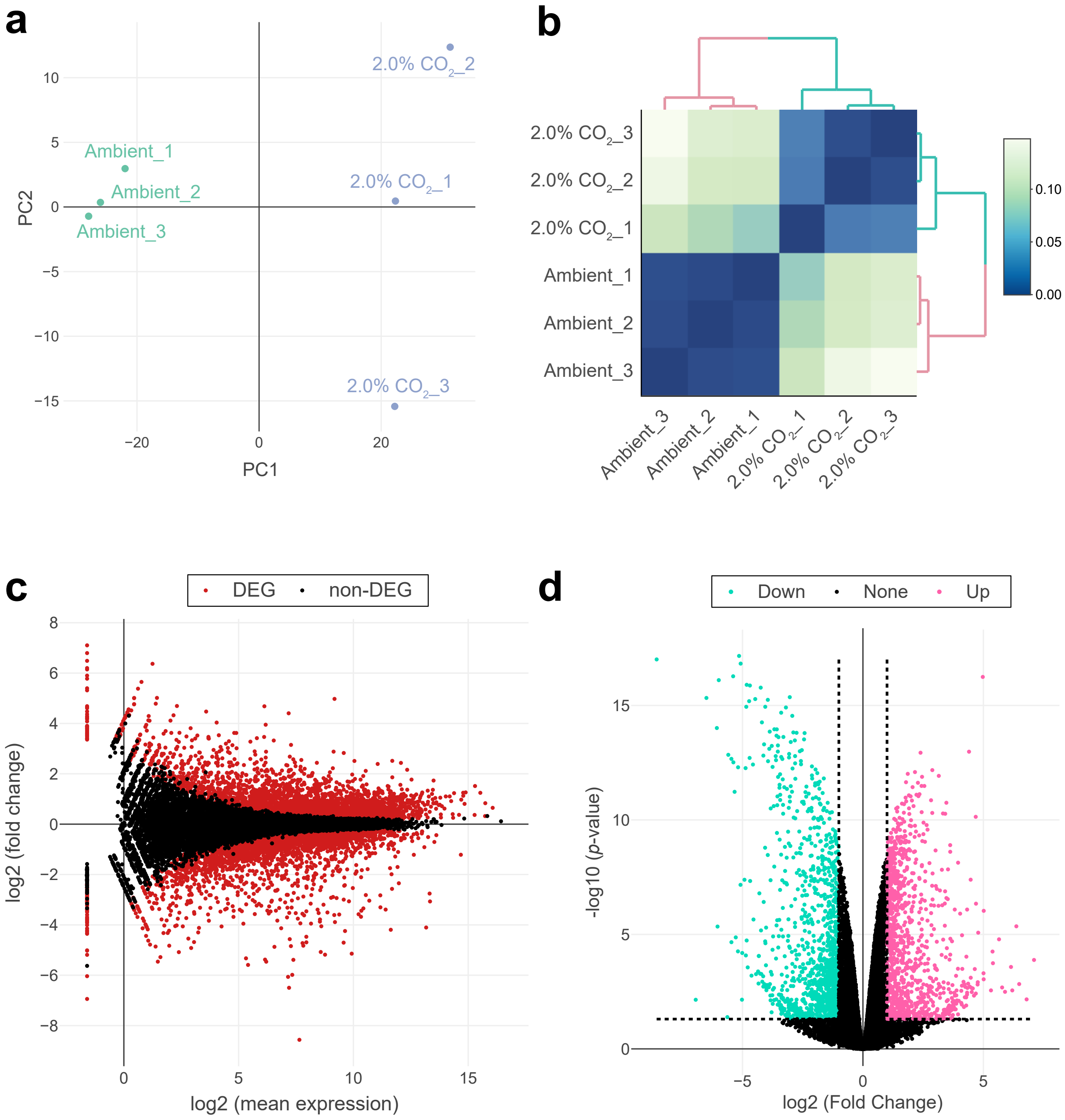
a
b
c
d
PC1
PC2
Ambient_1
Ambient_2
Ambient_3
2.0% CO2_1
2.0% CO2_2
2.0% CO2_3
DEG
non-DEG
log2 (mean expression)
log2 (fold change)
Down
None
Up
log2 (Fold Change)
-log10 (p-value)

**e**

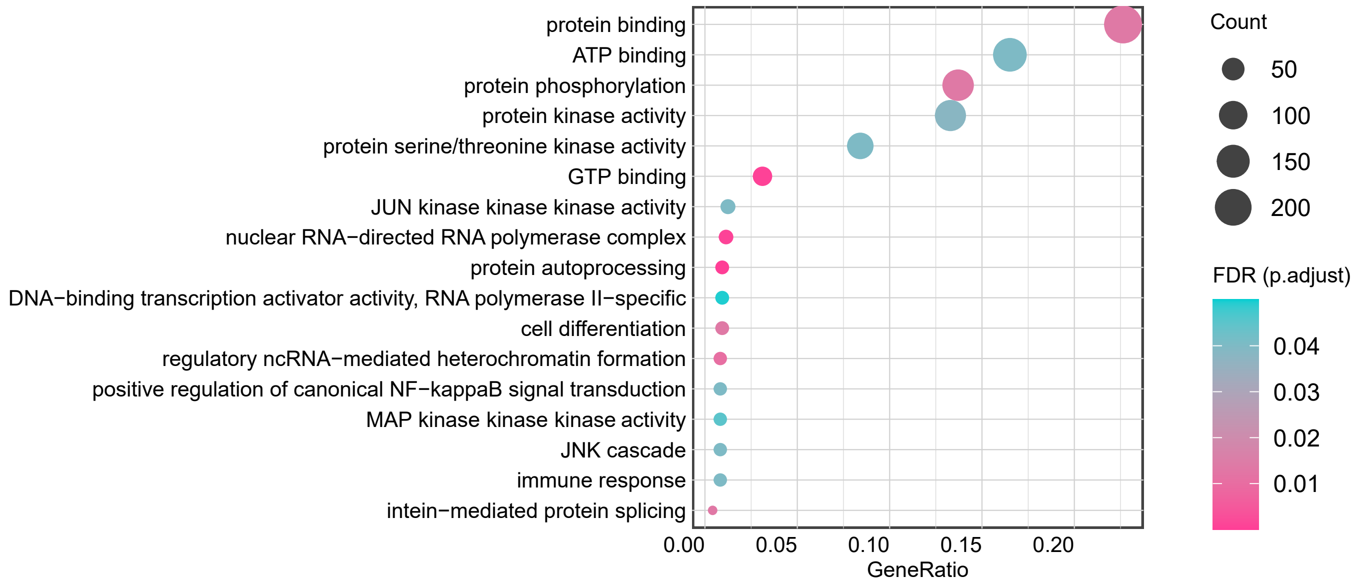
protein binding
ATP binding
protein phosphorylation
protein kinase activity
protein serine/threonine kinase activity
GTP binding
JUN kinase kinase kinase activity
nuclear RNA−directed RNA polymerase complex
protein autoprocessing
DNA−binding transcription activator activity, RNA polymerase II−specific
cell differentiation
regulatory ncRNA−mediated heterochromatin formation
positive regulation of canonical NF−kappaB signal transduction
MAP kinase kinase kinase activity
JNK cascade
immune response
intein−mediated protein splicing
0.00
0.05
0.10
0.15
0.20
GeneRatio
Count
50
100
150
200
FDR (p.adjust)
0.04
0.03
0.02
0.01

**f**

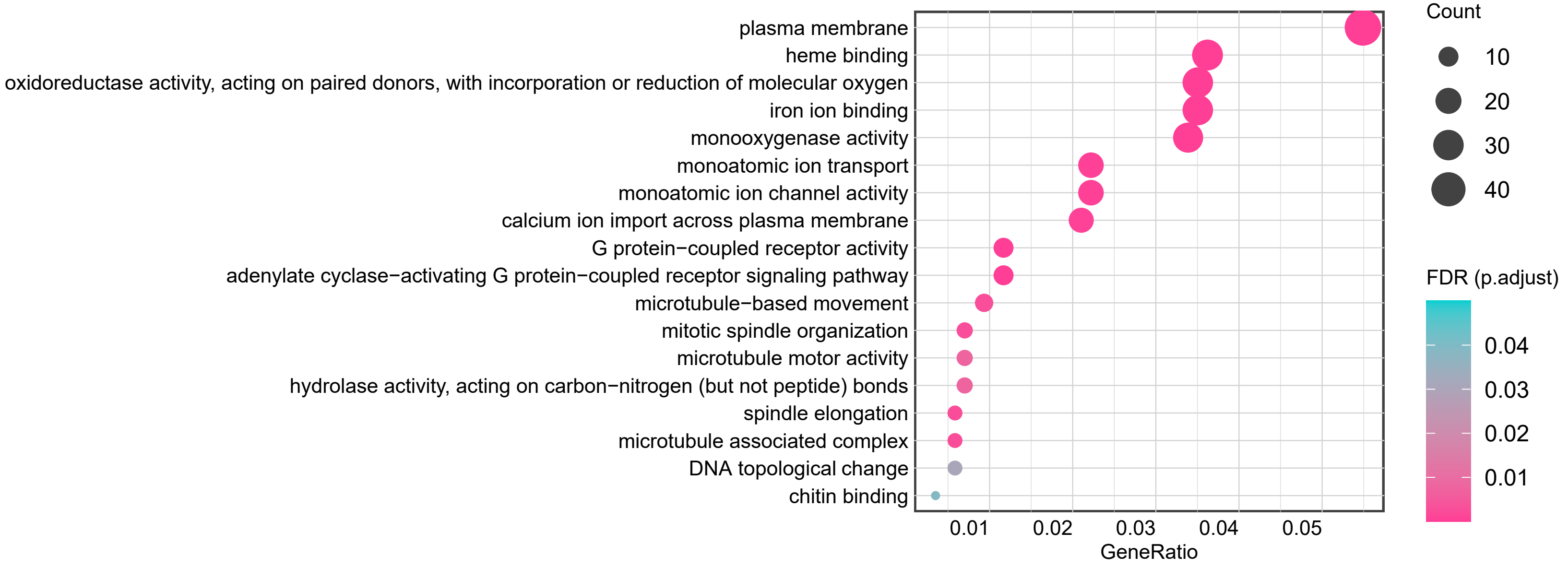
plasma membrane
heme binding
oxidoreductase activity, acting on paired donors, with incorporation or reduction of molecular oxygen
iron ion binding
monooxygenase activity
monoatomic ion transport
monoatomic ion channel activity
calcium ion import across plasma membrane
G protein−coupled receptor activity
adenylate cyclase−activating G protein−coupled receptor signaling pathway
microtubule−based movement
mitotic spindle organization
microtubule motor activity
hydrolase activity, acting on carbon−nitrogen (but not peptide) bonds
spindle elongation
microtubule associated complex
DNA topological change
chitin binding
0.01
0.02
0.03
0.04
0.05
GeneRatio
Count
10
20
30
40
FDR (p.adjust)
0.04
0.03
0.02
0.01

**Supplementary Figure 2.** Transcriptomic profiling and DEG of *R. clarus* under ambient and 2.0% $CO_2$ conditions.

**a**, Principal component analysis (PCA) of the gene expression profiles based on the top 1000 genes with the highest variance. **b**, Sample-to-sample distance heatmap and hierarchical clustering based on overall gene expression patterns. Clustering was performed using the complete linkage method with Pearson correlation-based distance. The color scale indicates the distance between samples, where dark blue represents higher similarity. **c**, MA plot showing the relationship between mean expression ($\log_2$) and fold change ($\log_2$) across all detected genes. Red dots represent differentially expressed genes (DEGs), and black dots represent non-DEGs. **d**, Volcano plot displaying the distribution of gene expression changes. The y-axis represents $\log_{10}$ (*p*-value), and the x-axis represents $\log_2$ (fold change). Green and pink dots indicate significantly downregulated and upregulated genes, respectively, under 2.0% $CO_2$ condition. Black dots represent genes that did not meet the significance threshold. satisfying the criteria of $|\ \log_2$ fold change $| > 1$ and FDR $< 0.05$. Black dots represent genes that did not meet these significance thresholds. **e–f**, Gene Ontology (GO) enrichment analysis of DEGs. Enriched GO terms for upregulated (**e**) and downregulated genes (**f**) under 2.0% $CO_2$ condition. Dot size indicates the number of DEGs associated with each term (Count), and dot color represents the significance level.

**Supplementary Data 1.** RNA-seq analysis of *R. clarus* under 2.0% $CO_2$ compared with ambient conditions.

This file contains quality control results for RNA-seq, differential expression statistics for all analysed genes, and lists of up- and down-regulated genes.
